# Microbial metabolism and adaptations in *Atribacteria*-dominated methane hydrate sediments

**DOI:** 10.1101/536078

**Authors:** Jennifer B. Glass, Piyush Ranjan, Cecilia B. Kretz, Brook L. Nunn, Abigail M. Johnson, Manlin Xu, James McManus, Frank J. Stewart

**Affiliations:** School of Earth and Atmospheric Sciences, Georgia Institute of Technology, Atlanta, GA, USA; School of Biological Sciences, Georgia Institute of Technology, Atlanta, GA, USA; Department of Genome Sciences, University of Washington, Seattle, WA; Bigelow Laboratory for Ocean Sciences, East Boothbay, ME, USA; Department of Microbiology & Immunology, Montana State University, Bozeman, MT, USA

## Abstract

Gas hydrates harbor gigatons of natural gas, yet their microbiomes remain understudied. We bioprospected 16S rRNA amplicons, metagenomes, and metaproteomes from methane hydrate-bearing sediments under Hydrate Ridge (offshore Oregon, USA, ODP Site 1244, 2-69 mbsf) for novel microbial metabolic and biosynthetic potential. *Atribacteria* sequences generally increased in relative sequence abundance with increasing sediment depth. Most Atribacteria ASVs belonged to JS-1-Genus 1 and clustered with other sequences from gas hydrate-bearing sediments. We recovered 21 metagenome-assembled genomic bins spanning three geochemical zones in the sediment core: the sulfate-methane transition zone, metal (iron/manganese) reduction zone, and gas hydrate stability zone. We found evidence for bacterial fermentation as a source of acetate for aceticlastic methanogenesis and as a driver of iron reduction in the metal reduction zone. In multiple zones, we identified a Ni-Fe hydrogenase-Na+/H+ antiporter supercomplex (Hun) in *Atribacteria* and *Firmicutes* bins and in other deep subsurface bacteria and cultured hyperthermophiles from the *Thermotogae* phylum. *Atribacteria* expressed tripartite ATP-independent (TRAP) transporters downstream from a novel regulator (AtiR). Atribacteria also possessed adaptations to survive extreme conditions (e.g., high salt brines, high pressure, and cold temperatures) including the ability to synthesize the osmolyte di-myo-inositol-phosphate as well as expression of K^+^-stimulated pyrophosphatase and capsule proteins.

**Originality-Significance Statement:** This work provides insights into the metabolism and adaptations of microbes that are ubiquitous and abundant in methane-rich ecosystems. Our findings suggest that bacterial fermentation is a source of acetate for aceticlastic methanogenesis and a driver of iron reduction in the metal reduction zone. *Atribacteria*, the most abundant phylum in gas hydrate-bearing sediments, possess multiple strategies to cope with environmental stress.

## Introduction

Gas clathrate hydrates are composed of solid water cages encasing gas molecules, commonly methane (CH_4_). Methane hydrates form naturally under high pressure and low temperature along continental margins (Kvenvolden, 1993; Mazurenko and Soloviev, 2003; Hester and Brewer, 2009; Collett et al., 2015). Continental margins and shelves harbor gigatons of natural gas in hydrates, which are susceptible to dissociation due to rising ocean temperatures, with potential for releasing massive methane reservoirs to the ocean and the atmosphere, which could exacerbate global warming (Archer et al., 2009; Ruppel and Kessler, 2017).

Despite the global importance of gas hydrates, their microbiomes remain largely unknown. Microbial cells are physically associated with hydrates (Lanoil et al., 2001), and the taxonomy of these hydrate-associated microbiomes is distinct from non-hydrate-bearing sites (Inagaki et al., 2006), possibly due to more extreme environmental conditions. Because salt ions are excluded during hydrate formation, porewaters of hydrate-bearing sediments are hypersaline (Ussler III and Paull, 2001; Bohrmann and Torres, 2006). Hydrate-associated microbes may possess adaptations to survive high salinity and low water activity, as well as low temperatures and high pressures (Honkalas et al., 2016). However, knowledge of the genetic basis of such adaptations is incomplete, as genomic data for hydrate communities are sparse and most hydrate microbiomes have been characterized primarily through single-gene taxonomic surveys.

Global 16S rRNA gene surveys show that the JS-1 sub-clade of the uncultivated bacterial candidate phylum *Atribacteria*, also known as *Caldiatribacteriota*, is the dominant taxon in gas hydrates (Reed et al., 2002; Inagaki et al., 2003; Kormas et al., 2003; Newberry et al., 2004; Webster et al., 2004; Inagaki et al., 2006; Webster et al., 2007; Fry et al., 2008; Kadnikov et al., 2012; Parkes et al., 2014; Yanagawa et al., 2014; Chernitsyna et al., 2016; Gründger et al., 2019) and in other marine and freshwater sediment ecosystems with abundant methane (Blazejak and Schippers, 2010; Gies et al., 2014; Carr et al., 2015; Hu et al., 2016; Nobu et al., 2016; Lee et al., 2018; Bird et al., 2019). The other major *Atribacteria* lineage, OP-9, primarily occurs in hot springs (Dodsworth et al., 2013; Rinke et al., 2013) and thermal bioreactors (Nobu et al., 2015). Marine *Atribacteria* are dispersed through ejection from submarine mud volcanoes (Hoshino et al., 2017; Ruff et al., 2019), and environmental heterogeneity may select for locally adapted genotypes. *Atribacteria* are highly enriched in anoxic, organic, and hydrocarbon rich sediments (Chakraborty et al., 2020; Hoshino et al., 2020) and have recently been discovered to be actively reproducing in the deep subsurface (Vuillemin et al., 2020). The phylogenetic diversity of *Atribacteria* genera suggests the potential for uncharacterized variation in functional niches.

*Atribacteria* appear to rely primarily on heterotrophic fermentative metabolisms. The high-temperature OP-9 lineage ferments sugars to hydrogen, acetate, and ethanol (Dodsworth et al., 2013; Katayama et al., 2020). The low-temperature JS-1 lineage ferments propionate to hydrogen, acetate, and ethanol (Nobu et al., 2016). Some JS-1 strains can also ferment short-chain n-alkanes (e.g. propane) into fatty acids by fumarate addition (Liu et al., 2019). Both JS-1 and OP-9 lineages possess genes encoding bacterial microcompartment shell proteins that may sequester toxic aldehydes and enable their condensation to carbohydrates (Nobu et al., 2016). Marine sediment JS-1 express genes to use allantoin as an energy source or chemical protectant and, unlike most deep subsurface bacteria, also encode a membrane-bound hydrogenase complex cotranscribed with an oxidoreductase, suggesting the ability for anaerobic respiration (Bird et al., 2019).

Here we examined the distribution, phylogeny, and metabolic potential of uncultivated JS-1 *Atribacteria* beneath Hydrate Ridge, off the coast of Oregon, USA, using a combination of 16S rRNA gene amplicon, metagenomic, and metaproteomic analysis. We found that *Atribacteria* from JS-1 Genus-1 are abundant throughout in the gas hydrate stability zone (GHSZ) and that they harbor numerous strategies for tolerance of osmotic stress, including many biosynthesis pathways for unusual osmolytes.

## Results and Discussion

### Geochemical gradients

Sediment core samples spanned three geochemical zones from 0-69 meters below seafloor (mbsf) at the ODP Site 1244 at Hydrate Ridge, off the coast of Oregon, USA (Fig. S1; Tréhu et al., 2003): the sulfate-methane transition zone (SMTZ; 2-9 mbsf; cores C1H2, C1H3, F2H4), the metal (iron/manganese) reduction zone (MRZ; 18-36 mbsf; cores F3H4, C3H4, E5H5), and the GHSZ (45-124 mbsf; cores E10H5, E19H5; Fig. 1A, Table S1). Sediment porewater methane concentrations (approximate, due to loss during sampling) increased from negligible at the seafloor to 8% by volume at 3-5 mbsf, and remained <5% below 5 mbsf, except for sample C3H4 (21 mbsf, MRZ) with ~18% methane (Fig. 1A). Sulfate dropped from 28 to <1 mM from 0-9 mbsf (the SMTZ) and remained <1 mM below 9 mbsf (Fig. 1A). In the MRZ, dissolved Mn and Fe peaked at 6 and 33 μM, respectively, while outside of the MRZ, dissolved Mn was ~1 μM and dissolved Fe was 7-20 μM (Fig. 1A). Dithionite-extractable (see Roy et al. (2013)), termed here “reactive”, Fe (0.4-1.4%) and Mn (0.002-0.012%) generally increased with depth (Table S1). Total organic carbon concentrations varied between 1-2 weight % (Table S1). Gas hydrate was observed from 45-125 mbsf in freshly recovered sediment cores, which contained up to 20% hydrate in the pore space, primarily as hydrate lenses or nodule patches (Tréhu et al., 2004; Fig. 1A). Estimated *in situ* salinity ranged from seawater salinity (35 g kg^−1^) to >100 g kg^−1^ and was highest in the GHSZ (Milkov et al., 2004). *In situ* temperature ranged from ~4°C at the seafloor to ~6-11°C in the GHSZ (ShipboardScientificParty, 2003).

**Figure 1:**
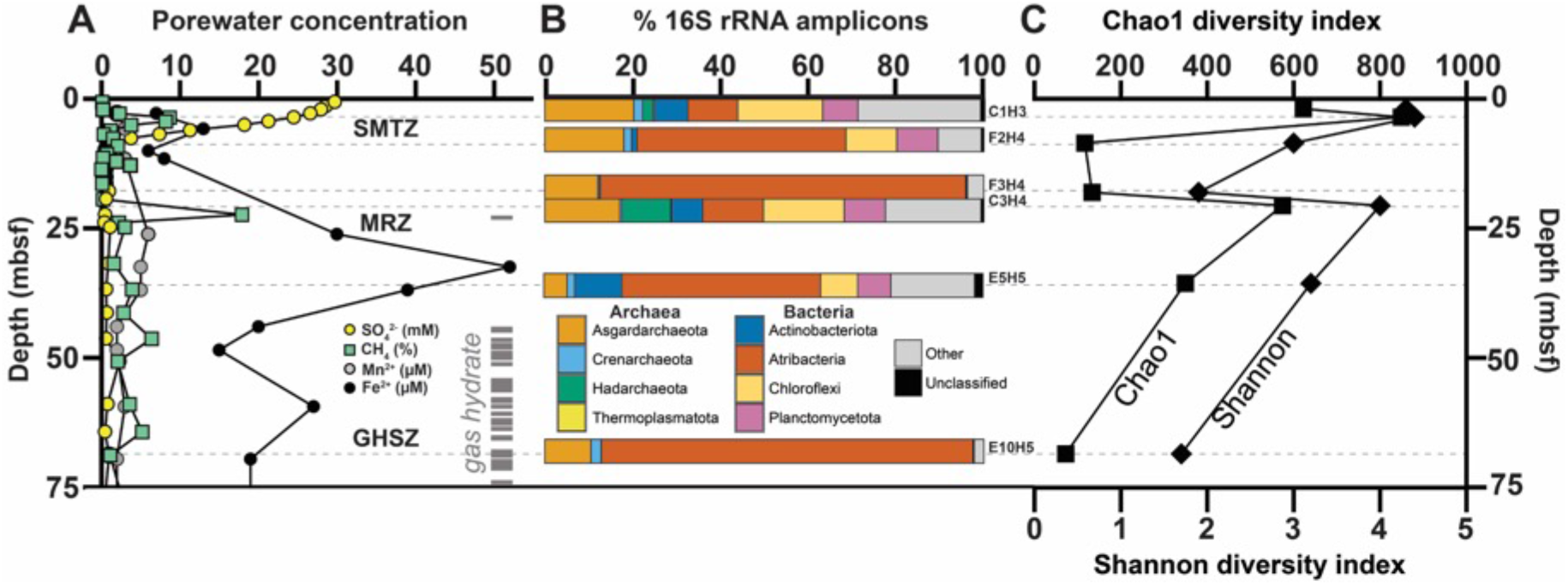
Porewater geochemistry and microbial taxonomy and diversity from sediment depth profiles at ODP 204 Site 1244, Hydrate Ridge, offshore Oregon, USA. **A:** Sulfate (yellow circles), methane (green squares), manganese (gray circles), and iron (black circles) concentrations, and depth of gas hydrate occurrences (gray dashes) from Tréhu et al. (2003). Sulfate and methane data are from core 1244B. Iron and manganese data are from core 1244E. Gas hydrate occurrence data are from core 1244C and 1244E. SMTZ: sulfate-methane transition zone; MRZ: metal reduction zone; GHSZ: gas hydrate stability zone. **B:** 16S rRNA gene amplicon taxonomic composition at the phylum level. “Other” category represents bacterial and archaeal phyla with <10% of total sequences. “Unclassified” represents sequences that were not classified at the phylum level. 16S rRNA amplicon data not shown for core C1H2 (see text). **C:** Microbial diversity based on Chao1 (top axis, squares) and Shannon index (bottom axis, diamonds) for the same 16S rRNA gene amplicon samples as shown in panel B.

### Atribacteria dominate ASVs in gas hydrate stability zone

*Actinobacteria, Atribacteria, Chloroflexi,* and *Planctomycetota* were the dominant bacterial phyla at Site 1244 (Fig. 1B). *Asgardarchaeota* and *Thermosplasmata* were the dominant archaeal phyla, with a notable rise in *Hadesarchaea* in the MRZ. Phylogenetic diversity in 16S rRNA gene amplicons based on the Shannon index and species richness based on the Chao1 index were highest in the SMTZ and MRZ, and lowest in the zones dominated by *Atribacteria*, in between the SMTZ and MRZ, and in the GHSZ (Fig. 1C). The relative sequence abundance of *Atribacteria* 16S rRNA amplicons ranged from 10-15% in the near surface to 80-85% at the top of the MRZ and the GHSZ (Fig. 1B). Most of the *Atribacteria* ASVs (n=20) belonged to JS-1 Genus 1 and clustered with other seep- and hydrate-associated sequences (Fig. 2A). ASV_368 comprised 54% of all amplicons in the GHSZ, and most GenBank sequences with 100% similarity to ASV_368 were from hydrate-bearing sediments from the Pacific Ocean basin (Table S2).

**Figure 2:**
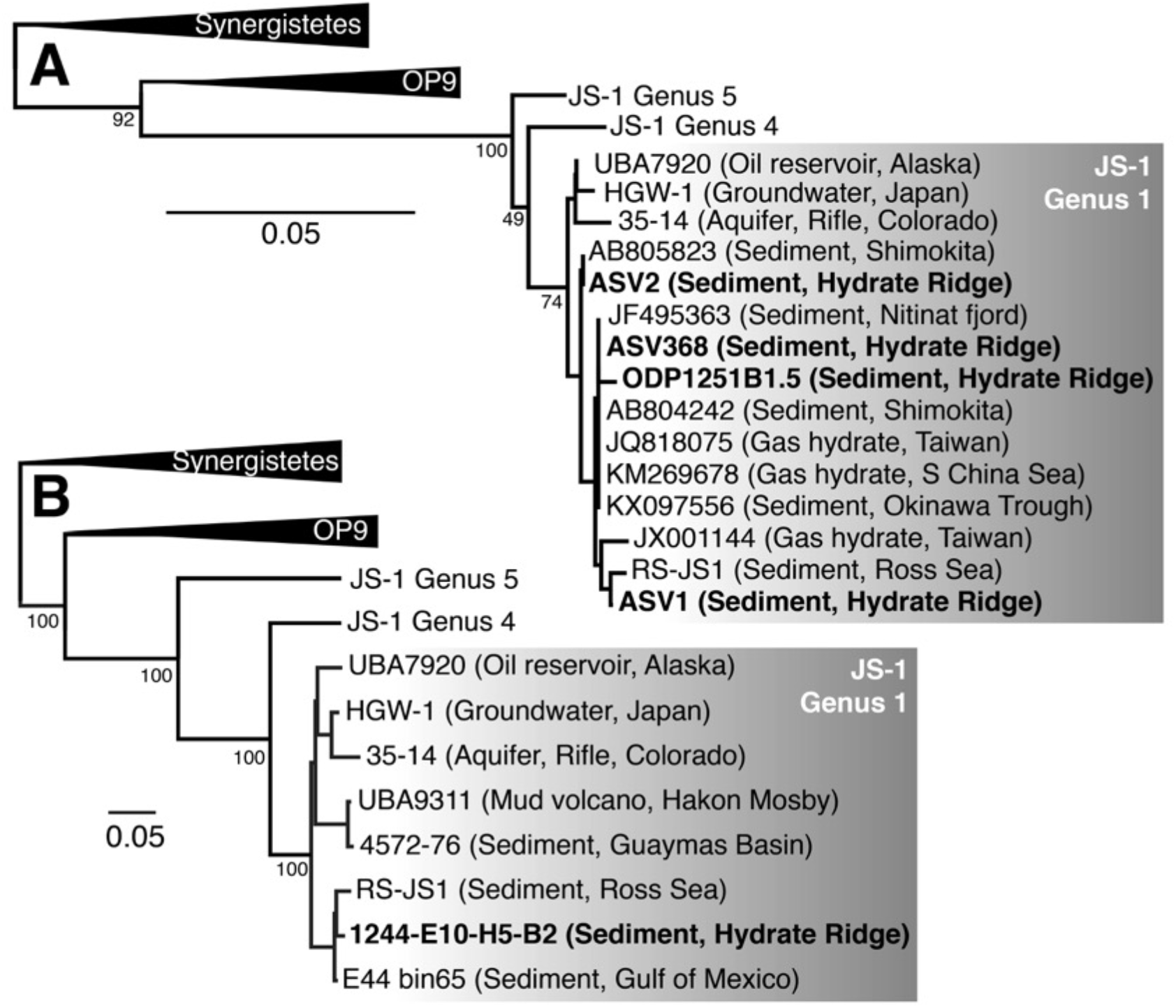
Neighbor-joining *Atribacteria* JS-1 Genus 1 phylogenies based on (A) 16S rRNA amplicons and (B) ribosomal proteins. Bolded sequences are from Hydrate Ridge. 16S rRNA phylogeny including the top three most abundant Atribacteria ASVs from Site 1244 (see Table S2 for relative sequence abundances). ODP1251B1.5 is the dominant JS-1 16S rRNA clone from Hydrate Ridge Leg 204 cores as reported by Inagaki et al. (2006). Italicized names are from MAGs or SAGs; the rest of the sequences are from 16S amplicons. Genera labels are based on sequences from Yarza et al. (2014) and Liu et al. (2019). Conserved sites used in phylogenies: (A) 190 bases; (B) 846 amino acids.

Metagenome-assembled binning yielded 21 MAGs with >35% completeness and <10% contamination including 17 bacteria and 4 archaea (Table S3). These MAGs included five *Dehalococcoidia* (*Chloroflexi*) in the SMTZ, MRZ, and GHSZ, and five *Firmicutes* (*Clostridia*) in the SMTZ and MRZ (Table S3). Other MAGs included one *Calditrichaeota* in the SMTZ, one each of *Bacteroidetes*, *Spirochaeta, Hadesarchaea*, and *Methanosarcinales* (*Euryarchaeota*) in the MRZ, and one *Atribacteria* in the GHSZ (MAG E10H5-B2). The higher relative sequence abundance of *Atribacteria* 16S rRNA sequences at the top of the MRZ and the GHSZ is consistent with higher read recruitment (~4-8%) of *Atribacteria* MAG E10H5-B2 metagenomes from those zones vs. other depths (<1%). Although E10H5-B2 lacked a 16S rRNA gene, the 16S rRNA gene in RS-JS1 was 99.68% identical to 16S rRNA sequences from hydrate sediments from offshore Japan (Shimokita Peninsula) and Taiwan (Lin et al., 2014), and 99.43% identical to a clone from the South China Sea (Li and Wang, 2013; Fig. 2A, Table S2). Phylogeny based on eight concatenated ribosomal proteins confirmed that MAG E10H5-B2 belonged to JS-1 Genus 1, and formed a monophyletic group with MAGs from petroleum seeps in the Gulf of Mexico (E44-bin65; Dong et al., 2019; Chakraborty et al., 2020) and marine sediments in the Ross Sea (RS-JS1; Lee et al., 2018; Fig. 2B). Average amino acid identities between MAGs in JS-1 Genus 1 was 72-83% (Table S4).

### Anaerobic hydrocarbon degradation, aceticlastic methanogenesis, and fermentative iron reduction

A recent study suggested that JS-1 can anaerobically degrade short-chain *n*-alkanes using fumarate addition enzymes (FAEs; Liu et al., 2019). Like oil reservoir *Atribacteria*, MAG E10H5-B2 contained genes encoding the glycyl radical protein subunit A (*faeA,* RXG63988, in the pyruvate formate lyase family) and D (*faeD*, RXG63989) on the same contig. However, the *faeA* gene product in MAG E10H5-B2 was shorter (786 aa) than in the oil reservoir MAGs (~860 aa) and the *fae* operon lacked the signature *faeC* gene between *faeD* and *faeA* that is characteristic of FAEs, suggesting that they may produce a different product.

Byproducts of fumarate addition enzymes (e.g., fatty acids) can be further degraded by other bacterial fermentation in marine sediments*. Firmicutes* degrade benzoate to acetate and transfer the electrons to crystalline Fe(III) minerals, producing dissolved Fe^2+^; thereafter, the acetate is converted to methane by syntrophic aceticlastic methanogenic archaea (Aromokeye et al., 2020). The relative sequence abundance of *Firmicutes* and the presence of aceticlastic *Methanosarcinales* (MAG F3H4-B6; Table S3) indicate that fermentative iron reduction was likely the source of the Fe^2+^ peak and the methane peak in the MRZ **(Fig. 1A)**. Acetate for aceticlastic methanogenesis could also come from other acetogens including *Atribacteria* (Carr et al., 2015).

A biogenic source of methane to the gas hydrates at Hydrate Ridge is consistent with previous isotopic analyses (Kastner et al., 1998). Production of acetate by fermentative bacteria below the SMTZ challenges the previous paradigm that acetate was completely consumed in the SMTZ and therefore that biogenic methane in gas hydrates originated solely from hydrogenotrophic methanogenesis via CO_2_ reduction (Whiticar et al., 1995). However, the carbon isotopic composition of methane in the gas hydrate at Hydrate Ridge is more consistent with a CO_2_ reduction pathway, and there may be additional deeper sources of hydrogenotrophic methane that mask the contribution from aceticlastic methanogenesis.

### Predicted respiratory function of novel Hun supercomplex

Two MAGs (*Atribacteria* E10H5-B2 and *Firmicutes* E5H5-B3) contained genes for a putative operon encoding a 16-subunit respiratory complex, hereafter designated Hun. The *hun* operon was also present in *Atribacteria* MAGs and SAGs from Baltic Sea sediments (Bird et al., 2019), in diverse deep biosphere bacterial MAGs (e.g. *Atribacteria, Omnitrophica, Elusimicrobia*, *Bacteriodetes* (Fig. 3A, Table S5)), and in hyperthermophilic bacterial isolates from the genus *Kosmotoga* (*Thermotogae* phylum; Fig. 3A). *Atribacteria hun* genes likely encode a complex of four protein modules that couple H^+^ and Na^+^ translocation to H_2_ production (Fig. 3B) based on their similarity to characterized proteins (Schut et al., 2016).

**Figure 3:**
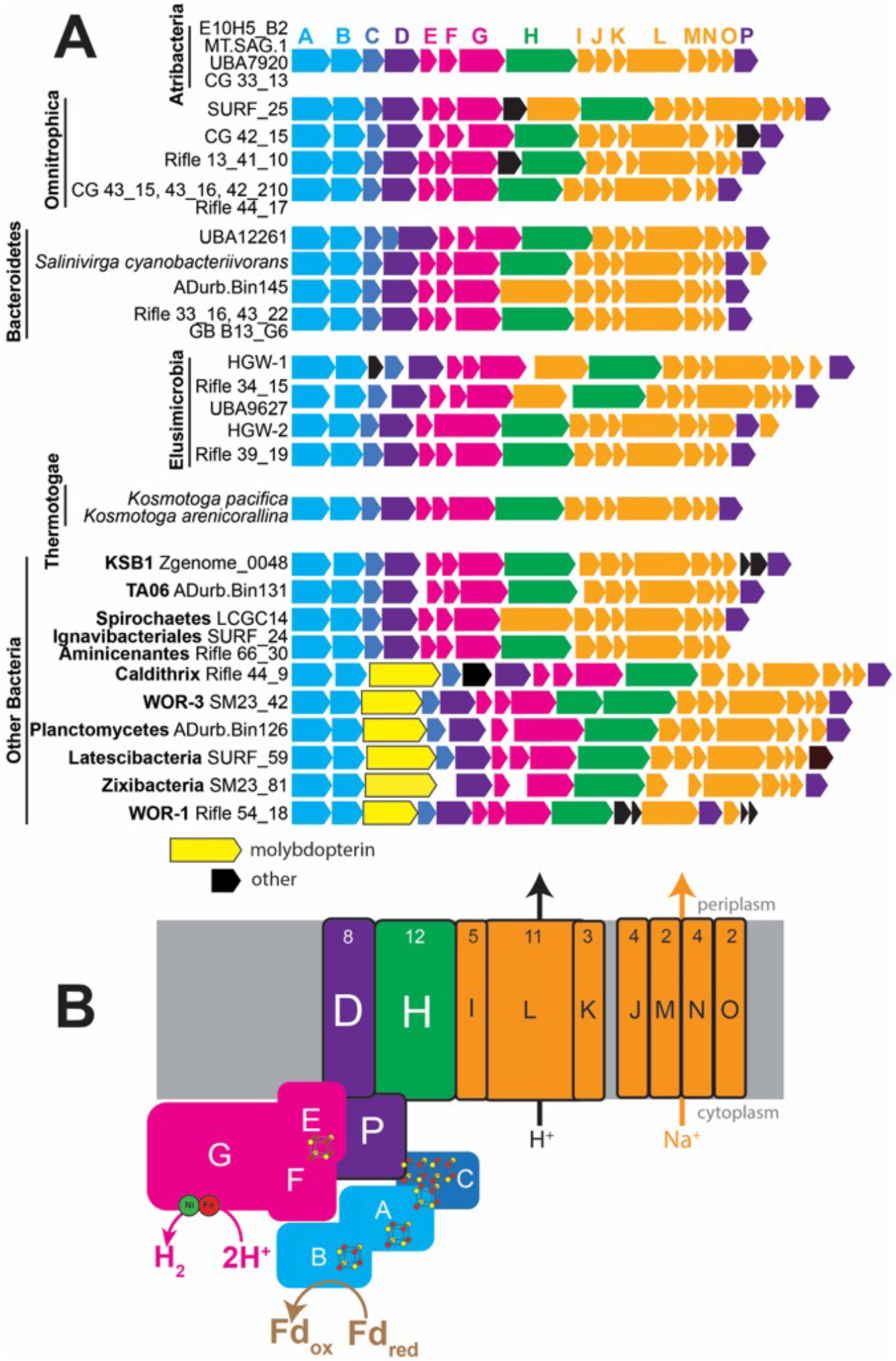
Gene neighborhood and predicted function of the predicted multi-subunit Hun respiratory complex. **A:** conserved gene cluster arrangement, with each color representing a different predicted protein. Some gene arrangements are found in more than one genome, as indicated. All MAGs and SAGs are from sediment samples. Sample abbreviations: ADurb: wastewater; CG: Crystal Geyser, Utah, USA; GB: Guaymas Basin, Gulf of California; HGW: Horonobe Underground Laboratory, Japan; LCGC: Loki’s Castle, Mid-Atlantic Ridge, Atlantic Ocean; MT: Mariana Trench; SM: White Oak Estuary, North Carolina, USA; SURF: Stanford Underground Research Facility, South Dakota, USA; Rifle: Rifle research site, Colorado, USA; UBA12261: wetland surface sediment; UBA9627: Rifle research site, Colorado, USA. **B:** predicted cellular locations and functions based on homologs of the genes of the same colors encoded by the putative *hun* operon in panel A. Iron-sulfur clusters and the Ni-Fe active site of HunG are also shown.

Additional analysis provided more insights into Hun function and phylogeny. The large hydrogenase subunit HunG was classified as a [NiFe] Group 4g-hydrogenase according to the Hydrogenase Database (Søndergaard et al., 2016). Group 4g-hydrogenases are biochemically unclassified but predicted to be ferredoxin-coupled and may couple reduced ferredoxin oxidation to proton reduction and H^+^/Na^+^ translocation (Greening et al., 2016). Based on the similarity of HunAB to anaerobic sulfite reductase (Asr) subunits A and B, which transfer electrons from ferredoxin to the active site in AsrC (missing in the *hun* operon), HunABC likely accept electrons from ferredoxin and pass them through iron-sulfur clusters to 2H^+^ for reduction to H_2_ at HunEFGP’s Ni-Fe active site, as suggested by the presence of two conserved CxxC motifs (L1 and L2) for Ni-Fe cofactor binding in HunG (Fig. 4A). Further analysis revealed that conserved residues for the Ni-Fe active site were different for HunG than other hydrogenases: CGIC-CYCC vs. CGIC-CxxC in other Group 4 hydrogenases (Fig. 4A). HunG was evolutionarily distant from other Group 4g sequences (Fig. 4A, B). In some MAGs, a 4Fe-4S molybdopterin domain-containing protein was present in between HunB and HunC (Fig. 3A). P-module-like subunits HunDHILK are predicted to be proton-pumping transmembrane proteins and Na-module-like subunits HunIJKLMNO are homologs of the Na^+^/H^+^ antiporter MnhABCDEFGH in Mrp-Mbh-type complexes. ATP is then generated via Na^+^-specific ATP synthases (Bird et al., 2019). Electrons from H_2_ could be transferred back to ferredoxin by the activity of the heterodisulfide reductase (HdrA)-methyl viologen hydrogenase (MvhAGD) complex.

**Figure 4:**
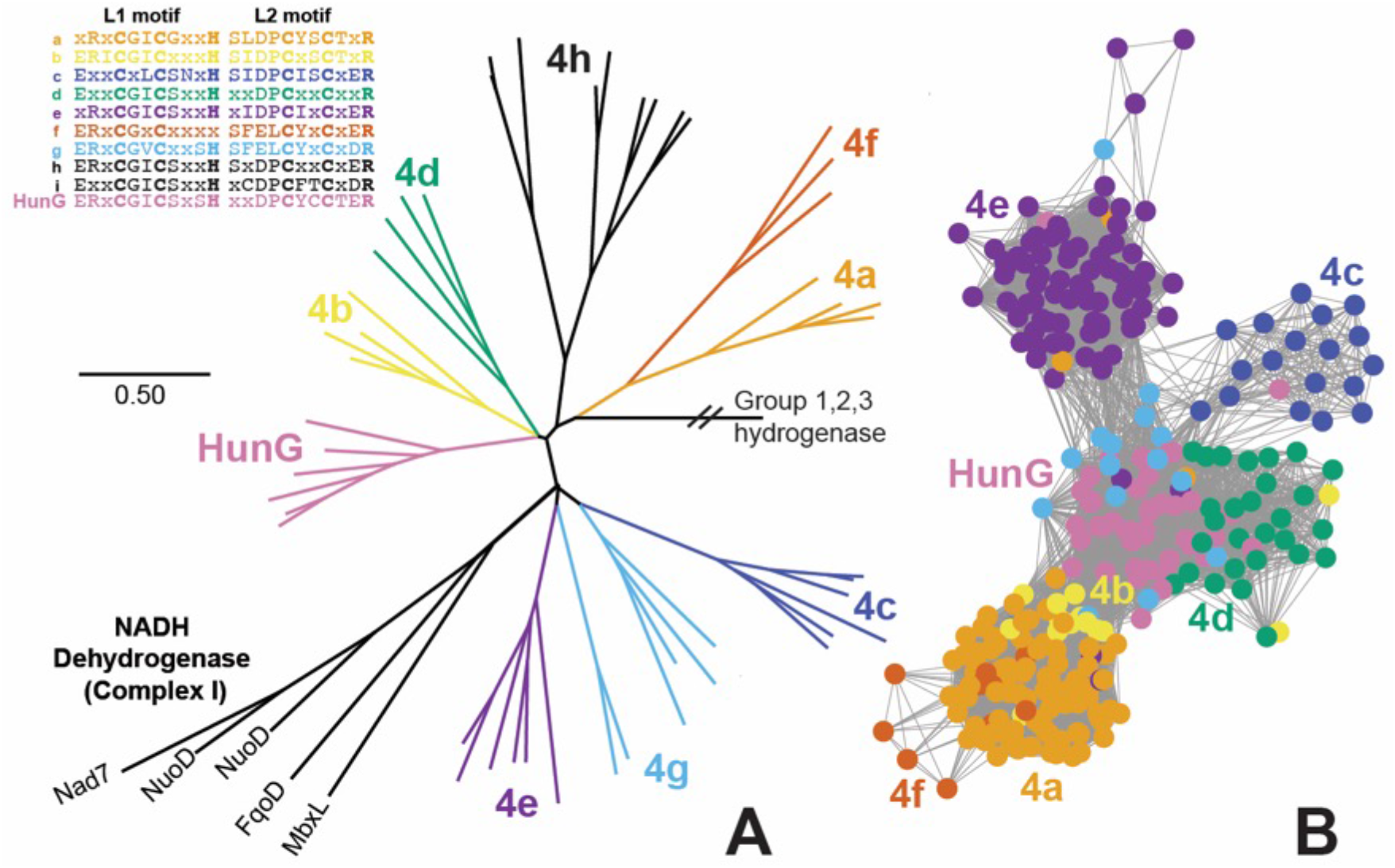
Phylogeny and sequence clustering of HunG and related large-subunit hydrogenases from group 4. **A:** Maximum likelihood HunG/NuoD/HycE phylogeny, with [Ni-Fe] hydrogenase group 4 labels drawn based on naming system from Søndergaard et al. (2016) and L1 and L2 motifs for the large subunit metal-binding centers for each class of group 4 hydrogenase. The NuoD subunit of NADH dehydrogenase (Complex I), which evolved from group 4 hydrogenase (Schut et al., 2016), is also included. **B:** Sequence similarity network for Group 4a,b,c,d,e,f,g and HunG hydrogenases with E-value cutoff of 10^−90^ and group color scheme the same as in A. Subgroups 4h and 4i are not shown in the sequence similarity network because they had no edges to the larger Group 4 cluster at E-value cutoff of 10^−90^.

### Transporters expressed in metaproteome

To assess gene expression, we analyzed metaproteomes from a subset of Site 1244 cores (C1H2, C3H4, and E10H5, from ~2, 20, and ~69 mbsf, respectively). Although we recovered few peptides of high quality, several robust hits were identified, including several types of transporters and cell envelope-associated proteins (Table 1). The expressed proteins were identified using *Atribacteria* MAGs from IODP Site 1244 as the reference database (see Methods) and had closest matches to other *Atribacteria* genomes (Table 1), suggesting that they originated from *Atribacteria*. Expressed proteins included a high-affinity branched-chain amino acid transport system permease (LivH) and multiple tripartite ATP-independent (TRAP) transporters (Fig. 5A). TRAP transporters use an electrochemical gradient (H^+^ or Na^+^) and a substrate-binding protein to transport a wide variety of molecules across the membrane (Rosa et al., 2018). Conserved residues within the TRAP substrate-binding protein confer specificity, with a conserved arginine residue essential for carboxylate transport (Fischer et al., 2015). In JS-1 MAGs from oil reservoirs, TRAP transport genes were associated with fumarate addition genes and likely transport fumarate or succinate for addition to hydrocarbons (Liu et al., 2019). In MAG E10H5-B2, genes for choline, inositol, and D-galacturonate catabolism often surrounded TRAP transporters (Fig. 5A), consistent with the finding that TRAP transporters can transport a much broader range of compounds than originally known (Vetting et al., 2015).

**Table 1.**
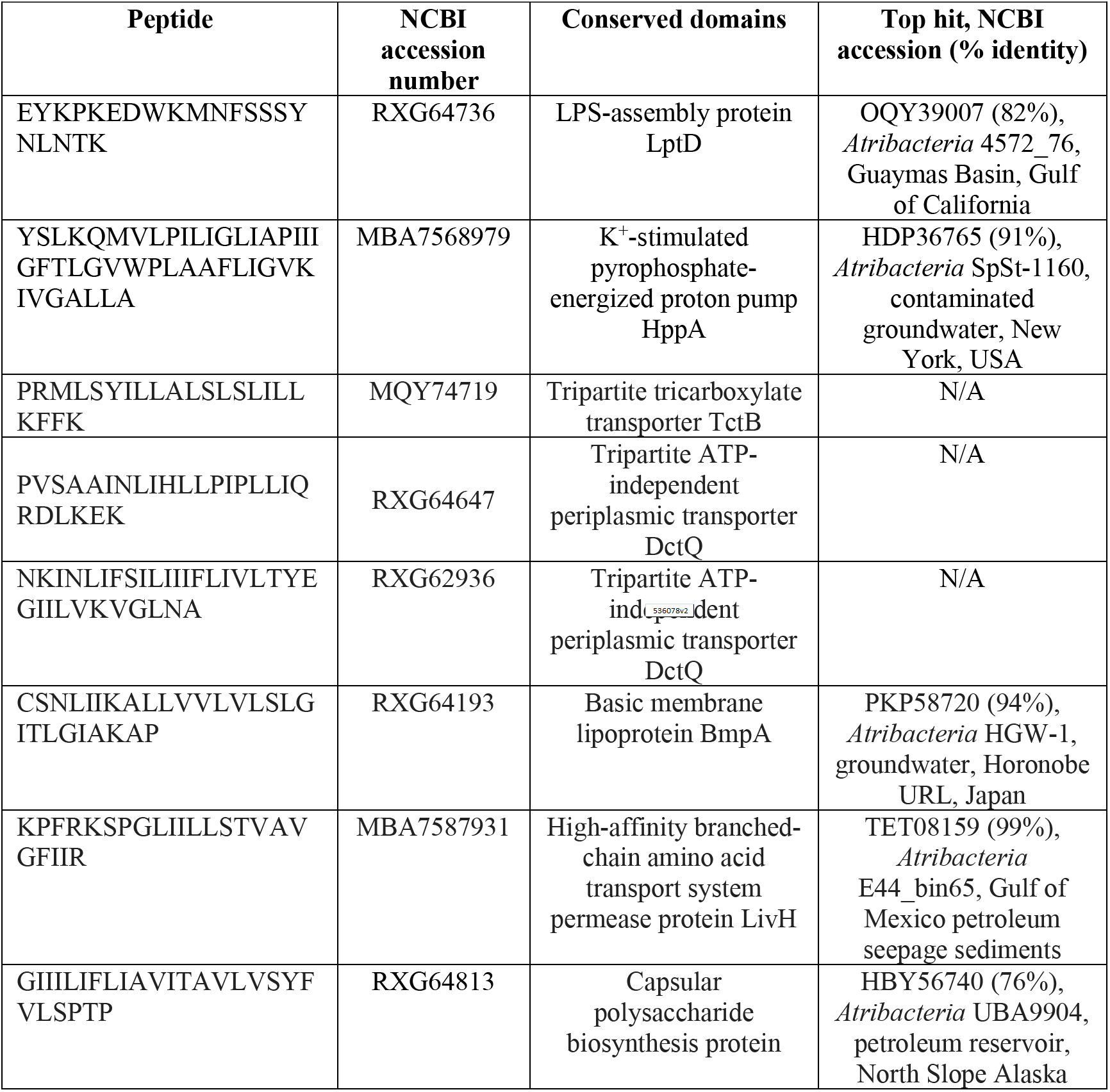
Peptide hits for ODP Site 1244 sample E10-H5 (~69 mbsf). Matches are shown for >70% identity to non-Hydrate Ridge genomes.

**Figure 5:**
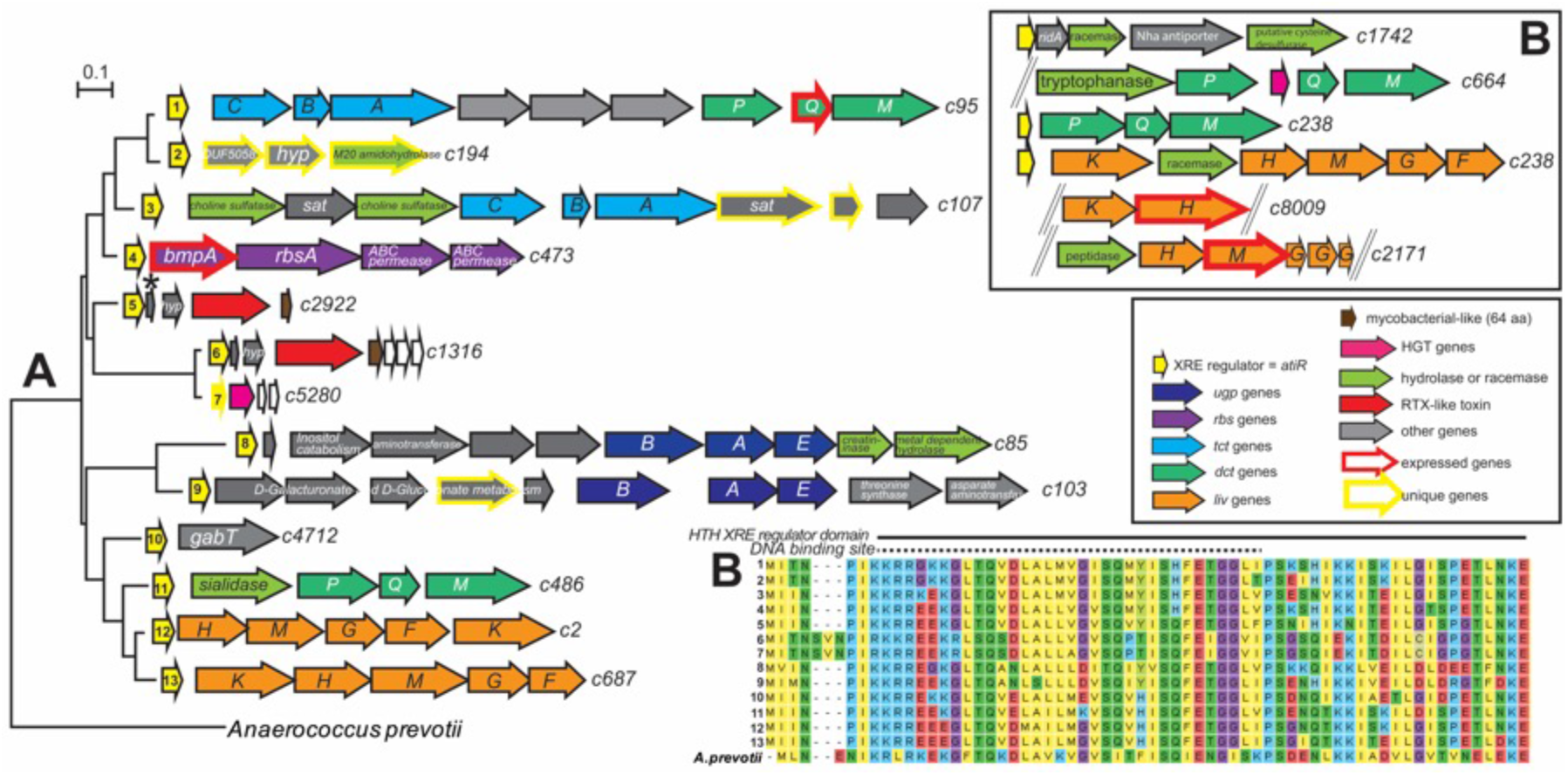
Phylogeny of helix-turn-helix xenobiotic response element regulators (yellow), hereafter “AtiR”, from B2 and synteny of downstream genes. Genes highlighted in thick red lines were expressed in the metaproteome. **A:** AtiR maximum likelihood phylogeny based on contigs (labeled on the right) from E10-H5 B2, with *Anaerococcus prevotii* as the outgroup. **Top inset:** Additional putative operons from B2 likely regulated by *atiR*, which is truncated partially or completely on these contigs. **Bottom inset:** Legend for panels A and B; **B:** AtiR amino acid alignment for the N-terminus of 13 AtiR sequences from *Atribacteria* E10-H5-B2 shown in panel A. Abbreviations: *bmpA*: basic membrane protein A; *dctPQM*: C4-dicarboxylate transporter; *gabT*: 4-aminobutyrate aminotransferase; *livHMGF*: branched chain amino acid transporter; *rbs*: ribose transporter; *sat:* sulfate adenylyltransferase; *tctCBA*: tricarboxylate transporter; *ugpBAE*: sn-glycerol-3-phosphate transporter. See Table S6 for accession numbers and % identity to closest gene hits in other genomes.

### AtiR, a novel regulator

Several of the expressed transporter proteins were encoded by genes downstream from a novel gene predicted to encode an ~85-amino acid helix-turn-helix xenobiotic response element (XRE) transcriptional regulator, which we named “AtiR” (Table S6; Fig. 5). AtiR was present in other genomes from Site 1244, in an *Atribacteria* MAG from marine hydrothermal sediment from Guaymas Basin (Zhou et al., 2020), and in unbinned contigs from marine hydrate-bearing sediments from offshore Shimokita Peninsula (Kawai et al., 2014 mbsf, core S12H4;). AtiR was also found in *Firmicutes* from other depths in Site 1244, including *Clostridia* MAG 1244-F3-H4-B2, *Firmicutes* MAG 1244-F2-H4-B10, and *Aminicenantes* MAG 1244-C3H4-B23. AtiR was also found in *Omnitrophica* genomes from Mid-Cayman Rise vent fluid plumes and in JS-1 genomes from Aarhus Bay, Denmark. Genes downstream of *atiR* were dominated by transporters for organic solutes (*tct, dct, ugp*), branched chain amino acids (*liv*), hydrolases (choline sulfatase, sialidase, tryptophanase, cysteine desulfurase), peptidases, racemases, and RTX-toxin (Tables S6; Fig. 5A). XRE regulators are widely distributed across the tree of life and regulate diverse metabolic functions and oxidative stress responses, typically as repressors that bind to DNA (Fig. 5B) to prevent transcription in the absence of a ligand. Methane-hydrate bacteria may use AtiR to regulate cellular degradation of peptides and proteins to amino acids, either for nutrient acquisition or for survival under environmental stress (Bergkessel et al., 2016).

### Osmotic stress survival

Any life that can persist in brine pockets within methane hydrate must contend with high salinity (up to ~3x that of seawater) and low water potential. We found a K^+^ stimulated pyrophosphatase, which is involved in salt stress in other bacteria (López-Marqués et al., 2004; Tsai et al., 2014), expressed the GHSZ sample (Table 1). *Atribacteria* MAG E10H5-B2 also contained numerous genes for the “salt out” survival strategy, in which osmotic pressure is maintained by exporting cations (Wood, 2015). Cation export systems included efflux systems, mechanosensitive ion channels, and Na^+^-H^+^ antiporters (Table 2). *Atribacteria* MAG E10H5-B2 also contained numerous MazEF toxin-antitoxin systems (Table S7), which are involved in translational control during stress response (Culviner and Laub, 2018).

**Table 2.** Putative osmotic stress-related genes in *Atribacteria* MAG E10-H5 B2. *indicates multiple copies.

| Annotation | Gene | Accession | Top hit (% identity) | *Atribacteria* MAG with top hit | Top hit environment and reference for metagenomes |
| --- | --- | --- | --- | --- | --- |
| Na<sup>+</sup> efflux | *natB* | RXG65900 | TET06401 (98%) | E44_bin65 | Gulf of Mexico petroleum seep sediments (Chakraborty et al., 2020) |
| Na<sup>+</sup> channel | DUF554 | RXG63559 | TET10447 (99%) |  |  |
| K<sup>+</sup> transport | *trkAH* * | RXG63511<br>RXG63512 | TET06940 (97%)<br>TET06939 (99%) |  |  |
| Mechanosensitive ion channel | *mscS* | RXG63036 | TET10003 (97%) |  |  |
| Glutamine synthetase | *glnA* | RXG65164 | TET08352 (99%) |  |  |
| Trehalose transporter | *sugAB* | RXG66833-RXG66834 | KUK55397- (94%)<br>KUK55398 (99%) | 34_128 | Oil reservoir, North Slope, Alaska (Hu et al., 2016) |
| Threonine/lysine efflux | *rhtB* | RXG66248 | KUK55393 (91%) |  |  |
| Na<sup>+</sup>/H<sup>+</sup> antiporter | *mrpEFGB* | RXG65834-<br>RXG65838 | TFB09297-<br>TFB09301 (91-95%) | MT.SAG.1 | Marianas Trench (Peoples et al., 2019) |
| Aromatic amino acid exporter | *yddG* * | RXG63201 | TFB08968 (91%) |  |  |
| Glutamate synthase | *gltD* | RXG66270 | PKP56573 (94%) | HGW-1 | Horonobe Underground Laboratory, Japan (Hernsdorf et al., 2017) |
| Proline racemase | *prdF* | RXG63210 | PKP58887 (92%) |  |  |
| Poly-gamma glutamate synthase | *pgsCBW* | RXG66317-RXG66319 | PKP60458-PKP60460 (~90%) |  |  |
| DIPP synthesis pathway | MIPS/IPCT-DIPPS* | RXG66889-RXG66888 | HBY57541-HBY57542 (~80%) | UBA9904 | Haakon Mosby mud volcano, Barents Sea (Niemann et al., 2006) |

A second salt survival strategy is import and/or biosynthesis of osmolytes, most often polar, water-soluble, and uncharged organic compounds and/or extracellular polymers. For example, glycine betaine is abundant in saline fluids from deep sediment basins (Daly et al., 2016). *Atribacteria* MAG E10H5-B2 contained genes for transport of trehalose and biosynthesis of the common osmolytes glutamine, glutamate, and poly-gamma-glutamate, all of which had homologs in other *Atribacteria* MAGs (Table 2). A capsular polysaccharide biosynthesis protein was among the handful of confident peptide hits (Table 1). *Atribacteria* transcripts for trehalose synthesis and transport were also present in other marine sediments (Bird et al., 2019). *Atribacteria* MAG E10H5-B2 also contained multiple copies of the aromatic amino acid exporter *yddG*, one of the most highly transcribed *Atribacteria* genes in other marine sediments (Bird et al., 2019). B2 and another *Atribacteria* MAG from a marine mud volcano (UBA9904) encoded myo-inositol-1 phosphate synthase (MIPS)/bifunctional IPC transferase and DIPP synthase (IPCT-DIPPS) for the unusual solute di-myo-inositol-phosphate (DIP; Table 2), which was previously only known to be made by hyperthermophiles (Santos and Da Costa, 2002).

The capacity for glycosylation may be another adaptation for survival of salt stress (Kho and Meredith, 2018). *Atribacteria* MAG E10H5-B2 and other *Atribacteria* encoded the non-mevalonate pathway for isoprenoid biosynthesis (*ispDEFGH*), exopolysaccharide synthesis proteins, numerous glycosyltransferases for transferring UDP- and GDP-linked sugars to a variety of substrates, and several proteins related to N-linked glycosylation (Table S8). Carbohydrate active enzymes are secreted by *Atribacteria* (Orsi et al., 2018) and may be involved in stress response. *Atribacteria* MAG E10H5-B2 also encoded genes for propionate catabolism and a bacterial microcompartment superlocus with 94-99% amino acid identity to a *Atribacteria* SAG from the Marianas Trench (Fig. S2), which is thought to be involved in sugar and aldehyde metabolism in *Atribacteria* (Axen et al., 2014; Nobu et al., 2016).

### Adaptations to life in methane hydrates

Microbes in the GHSZ in deep subsurface sediments appear to contain unique adaptations for survival in an extreme system with high salinity, high pressure, and low temperatures. Other probable environmental stress adaptations may include glycosylation and membrane modifications. It is also possible that these microbes can produce secondary metabolites that modify gas hydrate properties; we recently showed experimentally that recombinant *Chloroflexi* proteins from metagenomic sequences native to methane hydrate-bearing sediments alter the structure of clathrates (Johnson et al., 2020). More experiments are required to resolve the complex metabolic pathways and biosynthetic potential of life in methane hydrates, with important implications for stability of gas hydrates on our own planet (e.g. Snyder et al., 2020) and potential habitability and survival strategies of other planetary bodies in our solar system (Mousis et al., 2015; Kamata et al., 2019).

## Acknowledgments

We thank Vinayak Agarwal, Brett Baker, Jennifer Biddle, Jordan Bird, Anirban Chakraborty, Frederick Colwell, Sheng Dai, Xiyang Dong, Konstantinos Konstantinidis, Peter Girguis, Julie Huber, Casey Hubert, Raquel Lieberman, Karen Lloyd, Katie Marshall, Alejandra Prieto Davo, Brandi Reese, Claudia Remes, Emil Ruff, Anne Trehu, Despina Tsementzi, Paula Welander, Loren Williams, Jieying Wu, and Jenny Yang for helpful discussions; Phil Rumford and curatorial staff at the ODP Gulf Coast Repository for providing samples; and Shweta Biliya, Annie Hartwell, Janet Hatt, Mike Lee, Nastassia Patin, Ben Tully, and the Bioinformatics Virtual Coordination Network for technical assistance with sequencing and bioinformatic analysis. This research was funded by Center for Dark Energy Biosphere Investigations (C-DEBI) Small Research Grant to J.B.G. and C.B.K. (NSF OCE-0939564), NASA Exobiology grant to J.B.G. and F.J.S. (NNX14AJ87G), NSF Biology Oceanography grant to F.J.S and J.B.G. (NSF OCE-1558916), NASA Exobiology grant to J.B.G. (80NSSC19K0477), and a Georgia Tech Earth and Atmospheric Sciences Frontiers Postdoctoral Fellowship to C.B.K. Metaproteomic analysis by B.L.N. was partially supported by the University of Washington’s Proteomic Resource (UWPR95794). This is C-DEBI contribution [provided upon paper acceptance].

## Experimental Procedures

### Sample collection

Sediments were cored at ODP site 1244 (44°35.1784′N; 125°7.1902′W; 895 m water depth; **Fig. S1**) on the eastern flank of Hydrate Ridge ~3 km northeast of the southern summit on ODP Leg 204 in 2002 (Tréhu et al., 2003) and stored at −80°C at the ODP Gulf Coast Repository.

### Geochemistry

Data for dissolved methane, sulfate, manganese, iron, and iodide in sediment porewaters were obtained from Tréhu et al. (2003). Reactive iron and manganese were extracted from frozen sediments using the citrate-dithionite method (Roy et al., 2013) and measured by inductively coupled plasma optical emission spectrometer (Agilent Technologies 700 Series). Total carbon, total nitrogen and total sulfur were determined by CNS analyzer (Perkin Elmer 2400). Total inorganic carbon was measured by CO_2_ coulometer (CM5130) with a CM5130 acidification module. Geochemical metadata are given in Table S1 and archived in BCO-DMO project 626690.

### DNA extraction

DNA was extracted, in duplicate, from 8-20 g of sediment from the following depths in meters below seafloor (mbsf, using IODP core designations, see (ShipboardScientificParty, 2003)): 1.95-2.25 (C1-H2); 3.45-3.75 (C1-H3); 8.60 (F2-H4); 18.10 (F3-H4); 20.69 (C3-H4); 35.65 (E5-H5); 68.55 (E10-H5); 138.89 (core E19-H5) using a MO-BIO PowerSoil total RNA Isolation Kit with the DNA Elution Accessory Kit, following the manufacturer protocol without beads. DNA pellets from two replicates from each depth were pooled together. DNA concentrations were measured using a Qubit 2.0 fluorometer with dsDNA High Sensitivity reagents (Invitrogen, Grand Island, NY, USA). DNA yields ranged from 4-15 ng per gram of sediments. Core E19-H5 (139 mbsf) yielded only 2 ng DNA per gram of sediment and yielded unreliable data due to contamination with sequences from the enzymes used in the library preparations. Therefore, this core segment was excluded from further analysis.

### 16S rRNA gene amplicon sequencing

Microbial community composition was assessed by Illumina sequencing of the V3-V4 region of the 16S rRNA gene. The V3-V4 region was PCR-amplified using primers F515 and R806 (Caporaso et al., 2011), each appended with barcodes and Illumina-specific adapters according to (Kozich et al., 2013). Reactions consisted of 1-2 μL DNA template (2 ng), 5 μL of 10x Taq Mutant reaction buffer, 0.4 μL of Klentaq LA Taq Polymerase (DNA Polymerase Technology, St. Louis, MO, USA), 2 μL of 10 mM dNTP mix (Sigma Aldrich, St. Louis, MO, USA), 2 μL of reverse and forward primers (total concentration 0.4 μM), and DNA-free water (Ambion, Grand Island, NY, USA) for the remainder of the 50 μL total volume. PCR conditions were an initial 5-min denaturation at 94°C, followed by 35 cycles of denaturation at 94°C (40 sec), primer annealing at 55°C (40 sec), and primer extension at 68°C (30 sec). Amplicon libraries were purified using a QIAquick PCR Purification Kit (Qiagen, Germantown, MD, USA), quantified by Qubit (Life Technologies), and pooled in equimolar concentration. Amplicons were sequenced on an Illumina MiSeq across two runs using the V2 500-cycle kit with 5% PhiX to increase read diversity. 16S rRNA sequences were deposited into NCBI SAMN04214977-04214990 (PRJNA295201).

### 16S rRNA gene amplicon taxonomic analysis

16S rRNA sequences were trimmed using Trim Galore (criteria: length >100 bp length, Phred score >25). Sequences were dereplicated with a cutoff of 200 bp, chimeras were removed, and ASVs were resolved using deblur (Amir et al., 2017). Shannon and chao 1 diversity indices were calculated in R using phyloseq (McMurdie and Holmes, 2013).

### *Atribacteria* ASV phylogenetic analysis

The reference alignment included *Atribacteria* 16S rRNA gene sequences from environmental clones from Inagaki et al. (2006); Nobu et al. (2016), Carr et al. (2015), and Yarza et al. (2014). The alignment was trimmed to include only the V3-V4 region spanned by the *Atribacteria* ASV sequences, resulting in a final alignment with 198 bases. The DNA sequences were aligned in MAFFT with the L-INS-i option (Katoh and Standley, 2013). A neighbor-joining phylogeny with 100 bootstraps was rooted with members of the *Synergistetes* bacterial phylum.

### Multiple displacement amplification, library preparation, and sequencing

Genomic DNA was amplified from all samples using a REPLI-g Single Cell Kit (Qiagen, Germantown, MD, USA) using UV-treated sterile plasticware and reverse transcription-PCR grade water (Ambion, Grand Island, NY, USA). Quantitative PCR showed that the negative control began amplifying after 5 hr of incubation at 30°C, and therefore, the 30°C incubation step was shortened to 5 hr using a Bio-Rad C1000 Touch thermal cycler (Bio-Rad, Hercules, CA, USA). DNA concentrations were measured by Qubit. Two micrograms of MDA-amplified DNA were used to generate genome libraries using a TruSeq DNA PCR-Free Kit following the manufacturer’s protocol (Illumina, San Diego, CA, USA). The resulting libraries were sequenced using a Rapid-Run on an Illumina HiSeq 2500 to obtain 100 bp paired-end reads. Sequencing statistics are provided in Table S3.

### Metagenome assembly, binning, and annotation

Demultiplexed Illumina reads were mapped to known adapters using Bowtie2 in local mode to remove any reads with adapter contamination. Demultiplexed Illumina read pairs were quality trimmed with Trim Galore (Babraham Bioinformatics) using a base Phred33 score threshold of Q25 and a minimum length cutoff of 80 bp. Paired-end reads were then assembled into contigs using SPAdes assembler with --meta option for assembling metagenomes, iterating over a range of k-mer values (21,27,33,37,43,47,51,55,61,65,71,75,81,85,91,95). Assemblies were assessed with reports generated with QUAST. Features on contigs were predicted through the Prokka pipeline with Barrnap for rRNA, Aragorn for tRNA, Infernal and Rfam for other non-coding RNA, and Prodigal for protein coding genes. Metagenomic 16S rRNA sequences predicted by Barrnap were analyzed by BLASTN analysis against the SILVA SSU database version 138.

Annotation of protein-coding genes was performed as follows: 1) BLASTP search against the default set of core genomes, followed by HMM search against a set of default core HMM profiles available in Prokka, 2) use of the BLAST Descriptor Annotator algorithm in BLAST2GO, which conducts BLAST against the NCBI nr database, 3) KEGG orthology assignment using GhostKoala and 4) InterProScan analysis, which involves cross-reference HMM searches across multiple databases to find Pfam families with close homology. Metagenomic sequences were deposited into NCBI SAMN07256342-07256348 (PRJNA390944). Whole Genome Shotgun projects has been deposited at DDBJ/ENA/GenBank under the accession JABUBK000000000-JABUBQ000000000.

### Metagenome-assembled genomes

Metagenome contigs were partitioned through MetaBAT (Kang et al., 2015) into metagenome-assembled genomes (MAGs) using tetranucleotide frequency and sequencing depth. Sequencing depth was estimated by mapping reads on to assembled contigs using Bowtie2 and Samtools. Completeness, contamination and strain level heterogeneity were assessed using single copy marker genes in CheckM (Parks et al., 2015). Gene features and their functional annotations for genome bins were extracted from the metagenome for the contigs that belong to the bins. Taxonomic affiliation for each bin was inferred via the least common ancestor (LCA) algorithm in MEGAN6 and by the top BLAST matches to the marker gene *rpoB.* Twenty-one MAGs with estimated completeness >50% were deposited into GenBank (Table S3). The B2 MAG was deposited into GenBank as “Candidatus *Atribacteria* bacterium 1244-E10-H5-B2” (SAMN07342547; NMQN00000000.1). Read recruitments of metagenomic sequences to MAGs were performed using Bowtie2 (Langmead and Salzberg, 2012) normalized to the approximate number of genomes in the metagenome estimated with MicrobeCensus (Nayfach and Pollard, 2015). The average amino acid identity matrix was generated using the ANI-AAI matrix tool (Rodriguez-R and Konstantinidis, 2016).

### *Atribacteria* MAG and SAG phylogeny

Public *Atribacteria* single cell amplified genomes (SAGs) or MAGs (77 genomes, as of July 2020) were collected into a Genome Group workspace in Pathosystems Resource Integration Center (PATRIC; Wattam et al., 2014). Six ribosomal proteins from the large rRNA subunit (L2, L3, L4, L6, L16, L18) and two from the large rRNA subunit (S3 and S19) were collected from the *Atribacteria* genomes using the Features tab in Genome Group View. The eight ribosomal proteins were concatenated and the amino acid sequences were aligned in MAFFT with the L-INS-i option (Katoh and Standley, 2013). A neighbor-joining phylogeny with 1000 bootstraps was rooted with members of the *Synergistetes* bacterial phylum.

### Maximum likelihood phylogenies

Large subunit hydrogenase (HunG) and the xenobiotic response element regulator (AtiR) were made using sequences aligned in MAFFT with the L-INS-i option (Katoh and Standley, 2013). Neighbor-joining phylogenies were made with 100 bootstraps.

### Gene neighborhood diagrams

Gene neighborhood diagrams for the *hun* gene neighborhood were made using the gene neighborhood tool (GNT) in EFI web tools using a “single sequence BLAST” function in “Retrieve Neighborhood Diagrams” set to an E-value of 10^−5^ and a window size of 20 (Zallot et al., 2019). The input sequence was NCBI accession RXG63129 for HunG.

### Metaproteomic sample preparation, mass spectrometry, and data analyses

Proteins from E10-H5 were extracted from a 10 g of frozen sediment using a protocol adapted from Nicora et al. (2013). Briefly, 2.5 mL of desorption buffer (0.5 M NaCl, 0.1 M glycerol, 0.2% SDS, 6 M urea, 1 mM EDTA, 100 mM ammonium bicarbonate) and 2 mL of a pH-buffered amino acid solution (containing equimolar histidine, lysine, and arginine, all 83 g 1 L-1 in ultra-pure water, pH 7.0) was added to the sample on ice. The goal of the pH-buffered amino acid solution is to fill the electronegative mineral sites in the sample with positively charged amino acids to reduce absorption of proteins to the particles. Samples were vortexed 4x, alternating 5 minutes vortexing and 5 min ice. The sediment slurry was then sonicated with Bronson probe sonicator (4 x 30 s) to lyse cells and heated at 95°C for 5 min. The sediment was pelleted by centrifugation (10,000 x g, 30 min, 4°C), and the supernatant was collected and stored on ice. The sediment pellet was washed 2 more times with 3 mL desorption buffer and supernatants were combined. In order to remove the SDS prior to protein digestion and mass spectrometry analysis, the filter aided sample preparation (FASP) method was used (Ostasiewicz et al., 2010). Millipore Amicon 10 kDa filter units were used and cleaned following manufacturer’s directions. Samples were loaded on top of filters (~9 mL) and centrifuged (3000 rpm, 90 min, 4°C). To remove all SDS, proteins retained on the filter were rinsed 3 times by adding 5 mL of 8 M urea in 50 mM ammonium bicarbonate and repeating the prior centrifugation step. Iodoacetamide (3 mL, 15 mM) was added to samples, incubated in the dark at room temperature for 30 minutes, and then centrifuged (3000 rpm, 90 min, 4°C). Proteins were then rinsed two times with 10 mL of 100 mM ammonium bicarbonate and centrifuged to remove liquid (3000 rpm, 90 min, 4°C). To digest protein on the filter, 0.5 μg of trypsin (modified, sequencing grade, Promega) was added to the filter, topped with 2.5 mL of 25 mM ammonium bicarbonate, vortexed, and incubated 12 hr at room temperature. Filtrate was collected by centrifugation (3000 rpm, 90 min, 4°C), and SpeedVaced to near dryness at 4oC. Peptides were then resuspended in 50 μL of 2% acetonitrile and 0.1% formic acid and desalted using Nest Group C18 Proto centrifugal macro columns following manufacturer’s instructions. Each 10 μL sample was separated on a NanoAquity UPLC with a 60 min gradient (2-35% acetonitrile) and analyzed on a Thermo Scientific Orbitrap Fusion Tribrid Mass Spectrometer operated in top20 data dependent acquisition mode.

A protein database for identifying the collected fragmentation spectra was generated from *Atribacteria* MAGs (C1H2_C3H4ab_E10H5_contam.fasta). These databases were concatenated with 50 common contaminants, yielding a protein database of 10,325 proteins. To assign spectra to peptide sequences, correlative database searches were completed using Comet v. 2015.01 rev. 2 (Eng et al., 2013; Eng et al., 2015). Comet parameters included: trypsin enzyme specificity, semi-digested, allowance of 1 missed cleavage, 10 ppm mass tolerance, cysteine modification of 57 Da (resulting from the iodoacetamide) and modifications on methionine of 15.999 Da (oxidation). Minimum protein and peptide thresholds were set at *P* > 0.95 on Protein and Peptide Prophet (Nesvizhskii et al., 2003). Protein inferences from the whole-cell lysates were accepted by ProteinProphet if the thresholds noted above were passed, two or more peptides were identified, and at least one terminus was tryptic (Keller et al., 2002; Nesvizhskii et al., 2003; Pedrioli, 2010). For each peptide discussed in the manuscript, manual inspection of the spectral identification was completed. The mass spectrometry proteomics data have been deposited to the ProteomeXchange Consortium via the PRIDE partner repository (Vizcaíno et al., 2015) with the dataset identifier PXD012479 (https://www.ebi.ac.uk/pride/archive/ Login: Password: BP2V3yGA).

