## Supplemental Material for "Microbial metabolism and adaptations in *Atribacteria*-dominated methane hydrate sediments"

[#]Now at: Michigan Medicine, University of Michigan, Ann Arbor, Michigan, USA;

[§]Now at: Association of Public Health Laboratories, Manager of Emerging and Zoonotic Infectious Diseases, Silver Spring, MD, USA

**Running Title:** *Atribacteria* adaptions in methane hydrate ecosystem

**Dedication:** To Katrina Edwards

**Table S1**. Additional geochemical data for ODP Site 1244 at Hydrate Ridge drilled on IODP Leg 204. Note: data not available (ND) for sample E5H5 (35.65 mbsf). SMTZ: sulfate-methane transition zone; MRZ: metal reduction zone; GHSZ: gas hydrate stability zone. DNA sequencing from E19H5 was unsuccessful. TC: total carbon; TN: total nitrogen; TIC: total inorganic carbon; TOC: total organic carbon; C:N carbon:nitrogen.

| Hole | Zone | Depth (mbsf) | %TC | %TN | %TS | %TIC | %TOC | C:N | %$CaCO_3$ | reactive Fe (%) | reactive Mn (%) |
|---|---|---|---|---|---|---|---|---|---|---|---|
| C1H2 | SMTZ | 1.95/2.25 | 2.07 | 0.20 | 0.42 | 0.17 | 1.9 | 11.05 | 1.46 | 0.38 | 0.002 |
| C1H3 | SMTZ | 3.45/3.75 | 1.88 | 0.14 | 0.55 | 0.70 | 1.2 | 9.85 | 5.81 | 0.45 | 0.004 |
| F2H4 | SMTZ | 8.6 | 1.54 | 0.17 | 0.39 | 0.44 | 1.1 | 7.55 | 3.66 | 0.57 | 0.005 |
| F3H4 | MRZ | 18.1 | 1.55 | 0.24 | 0.64 | 0.08 | 1.5 | 7.14 | 0.68 | 0.80 | 0.004 |
| C3H4 | MRZ | 20.69 | 1.22 | 0.18 | 0.22 | 0.08 | 1.1 | 7.40 | 0.65 | 1.10 | 0.004 |
| E5H5 | MRZ | 35.65 | NA | NA | NA | NA | NA | NA | NA | NA | NA |
| E10H5 | GHSZ | 68.55 | 1.71 | 0.22 | 0.42 | 0.13 | 1.6 | 8.38 | 1.08 | 0.51 | 0.003 |
| E19H5 | GHSZ | 138.89 | 1.42 | 0.20 | 0.07 | 0.51 | 0.9 | 5.32 | 4.23 | 1.41 | 0.012 |

**Table S2. Atribacteria 16S rRNA ASVs in Site 1244 and locations with hits with 100% identity.**

| | Western Pacific | | | | | | | | | | | | Eastern Pacific | | | | | | | | |
|---|---|---|---|---|---|---|---|---|---|---|---|---|---|---|---|---|---|---|---|---|---|
| | Offshore Shimokita Peninsula | Nankai Trough ODP Site 1176A | Ogasawara Trench | Okinawa Trough | Eastern Japan Sea | Japan Trench | Shiribeshi Trough | Sea of Okhotsk | Taiwan gas hydrate | Maluku Strait Indonesia | South China Sea gas hydrate | Xisha Trough, South China Sea | Guaymas Basin | Hydrate Ridge | Anoxic fjord | Santa Monica basin | Santa Barbara basin | Costa Rica mud volcano | Juan de Fuca segment seafloor | Eastern Equatorial Pacific ODP Site 1226 | Peru Margin ODP Site 1240 |
| **Sample Type (S: sediment, W: water)** | S | S | S | S | S | W | S | S | S | S | S | S | S | S | S | S | S | S | S | S | S |
| ASV_1 | ■ | ■ | | | | | | ■ | ■ | | ■ | | | | | | | | ■ | ■ | ■ |
| ASV_2 | ■ | | ■ | ■ | ■ | ■ | ■ | ■ | ■ | | ■ | | | ■ | ■ | | | | ■ | | ■ |
| ASV_37 | ■ | ■ | | ■ | | ■ | | ■ | ■ | ■ | ■ | ■ | ■ | | | | | | | ■ | |
| ASV_364 | | | | | | | | | ■ | | ■ | | | | | | | | | | |
| ASV_368 | ■ | | | ■ | ■ | ■ | | | ■ | | ■ | | ■ | | | | | | | ■ | |
| ASV_374 | ■ | | | | ■ | | | | | | | | ■ | ■ | ■ | ■ | | ■ | | | |
| ASV_379 | ■ | | | | | | | | ■ | | | | | ■ | ■ | | | | | | |
| ASV_394 | ■ | ■ | | | ■ | | | | ■ | | | | | | | | | | | ■ | |
| ASV_428 | ■ | | | | | | | | ■ | | ■ | | | ■ | | ■ | | | | | |
| ASV_543 | ■ | | | | | | | | ■ | | | | | | | | | | | | |
| ASV_602 | | | | | | | | | ■ | | | | | | | | | | | | |
| ASV_1059 | ■ | | | | | | | | | | | | | | | | | | | | |
| ASV_1519 | ■ | | | | | | | | ■ | | | | | | | | | | | | |
| ASV_1521 | | | | | | | | | | | | | | | | | | | | | |
| ASV_1522 | ■ | | | | | | | | ■ | | | | | | | | | | | | |
| ASV_1525 | ■ | | | | | | | | | | | | | | | | | | | | |
| ASV_1602 | ■ | | | ■ | ■ | | | ■ | ■ | | | | ■ | ■ | ■ | | ■ | | | | |
| ASV_1610 | ■ | ■ | | | ■ | ■ | | | | | ■ | | | ■ | | | | | | | |
| ASV_1612 | ■ | ■ | | | | ■ | | | | | ■ | | | | ■ | | | | | | |
| ASV_1784 | | | | | | | | | | | | | | | | | | | | | |

| | Southern Ocean | | Atlantic Ocean | | | | | | Other | | | | | | |
|---|---|---|---|---|---|---|---|---|---|---|---|---|---|---|---|
| | Ross Sea, Antarctica | Antarctic continental shelf | Haakon Mosby mud volcano | Gabon continental margin | Gulf of Mexico cold seep | Tidal flat Wadden Sea | Aarhus Bay Denmark | Gulf of Cadiz mud volcano | Kazan mud volcano Mediterranean Sea | Amsterdam mud volcano Mediterranean Se | Simeulue forearc basin Indian Ocean | Trikala City Greece drinking water | Japan groundwater | Spitsbergen permafrost soil | Alaska petroleum reservoir |
| **Sample Type (S: sediment, W: water)** | S | S | S | S | S | S | S | S | S | S | S | S | S | S | W |
| ASV_1 | ■ | | | | ■ | | | | | | | | | | |
| ASV_2 | | | | ■ | | | | | | | ■ | | | | |
| ASV_37 | | | | | | | | | | | | | | | |
| ASV_364 | | | | | | | | | | | | | | | |
| ASV_368 | | | | | | | | | | | | | | | |
| ASV_374 | | | ■ | | ■ | | | ■ | | | | | | | |
| ASV_379 | | | | | | | | | | | | | | | |
| ASV_394 | | | | | | | | | | | | | | | |
| ASV_428 | | ■ | | | | | | | | | ■ | | | | |
| ASV_543 | | | | | | | | | | | | | | | |
| ASV_602 | | | | | | | | | | | | | | | |
| ASV_1059 | | | | | | | | | | | | | | | |
| ASV_1519 | | | | | | | | | ■ | | | | ■ | | |
| ASV_1521 | | | | | | | | | | | | | | | |
| ASV_1522 | ■ | | | | | | | | | | | | | | |
| ASV_1525 | | | | | | | | | | | | | | | |
| ASV_1602 | | | ■ | ■ | ■ | ■ | ■ | ■ | ■ | ■ | ■ | ■ | | ■ | ■ |
| ASV_1610 | | | | | | | | | | | | | | | |
| ASV_1612 | | | | | | | | | | | | | | | |
| ASV_1784 | | | | | | | | | | | | | | | |

| | Relative abundance (% Atribacteria / % all 16S rRNA amplicons) | | | | | | |
|---|---|---|---|---|---|---|---|
| | *near surface* | *sulfate-methane transition zone* | | *metal reduction zone* | | | *gas hydrate stability zone* |
| **Sample Type (S: sediment, W: water)** | **C1H2** | **C1H3** | **F2H4** | **F3H4** | **C3H4** | **E5H5** | **E10H5** |
| ASV_1 | | 6/1 | 4/2 | 6/5 | 35/5 | 58/28 | 8/7 |
| ASV_2 | 11/1 | 93/10 | 40/20 | 42/34 | 12/2 | 41/20 | 7/6 |
| ASV_37 | 8/1 | | | | 22/3 | 1/0 | 8/7 |
| ASV_364 | 36/5 | | | | 1/0 | | |
| ASV_368 | 23/3 | | | | | | 71/54 |
| ASV_374 | 14/2 | | | | | | |
| ASV_379 | | | | | | | |
| ASV_394 | 6/1 | | | | 8/1 | | |
| ASV_428 | 2/0 | | | | | | 4/3 |
| ASV_543 | 1/0 | | | | | | |
| ASV_602 | | | | | | | |
| ASV_1059 | | | | | | | |
| ASV_1519 | | | 37/19 | | | | |
| ASV_1521 | | | 9/5 | | | | |
| ASV_1522 | | | 7/3 | | | | 1/1 |
| ASV_1525 | | | 1/0 | | | | |
| ASV_1602 | | | | | 14/2 | | |
| ASV_1610 | | | | | 4/1 | | |
| ASV_1612 | | | 2/1 | | 4/1 | | |
| ASV_1784 | | | | | | | |

**Table S3. Quality evaluation and accession numbers of the 21 metagenome-assembled genomes from IODP Site 1244 generated in this study with >35% completeness and <10% contamination.**

| Assembly | WGS | BioSample | MAG ID | Taxon | Size (bp) | Contigs | CDS | N50 | GC (%) | Completeness (%) | Contamination (%) | Heterogeneity (%) |
|---|---|---|---|---|---|---|---|---|---|---|---|---|
| GCA_009619245 | WJOO00000000 | SAMN13181020 | 1244-C1-H3-B20 | Dehalococcoidia (Chloroflexi) | 1,744,267 | 338 | 1772 | 6,049 | 52 | 75 | 9 | 0 |
| GCA_009619075 | WJOY00000000 | SAMN13181030 | 1244-C3-H4-B3 | | 935,948 | 47 | 910 | 30,310 | 45 | 74 | 3 | 67 |
| GCA_009619025 | WJPC00000000 | SAMN13181034 | 1244-E5-H5-B4 | | 620,966 | 90 | 634 | 7,554 | 55 | 49 | 0 | 0 |
| GCA_009619235 | WJPE00000000 | SAMN13181036 | 1244-E5-H5-B9 | | 1,315,519 | 215 | 1385 | 7,652 | 53 | 71 | 5 | 75 |
| GCA_009619215 | WJPF00000000 | SAMN13181037 | 1244-E10-H5-B3 | | 971,417 | 200 | 1038 | 6,335 | 56 | 72 | 5 | 29 |
| GCA_009618505 | WJOQ00000000 | SAMN13181022 | 1244-F2-H4-B2 | Clostridia (Firmicutes) | 2,136,034 | 406 | 2041 | 6,910 | 40 | 79 | 4 | 50 |
| GCA_009619115 | WJOR00000000 | SAMN13181023 | 1244-F2-H4-B10 | | 974,022 | 276 | 956 | 3,720 | 43 | 39 | 0 | 0 |
| GCA_009619015 | WJOU00000000 | SAMN13181026 | 1244-F3-H4-B2 | | 2,140,048 | 47 | 2077 | 131,274 | 34 | 81 | 0 | 0 |
| GCA_009619095 | WJOT00000000 | SAMN13181025 | 1244-F3-H4-B3 | | 1,988,642 | 132 | 1758 | 24,484 | 41 | 90 | 5 | 33 |
| GCA_009618495 | WJPA00000000 | SAMN13181032 | 1244-C3-H4-B5 | | 998,110 | 85 | 881 | 18,112 | 42 | 57 | 2 | 0 |
| GCA_003575245 | NMQN00000000 | SAMN07342547 | 1244-E10-H5-B2 | Atribacteria | 4,055,260 | 912 | 4254 | 5,515 | 33 | 69 | 2 | 100 |
| GCA_009619225 | WJOX00000000 | SAMN13181029 | 1244-C3-H4-B19 | Bacteroidetes | 3,577,859 | 750 | 2889 | 6,011 | 38 | 77 | 6 | 14 |
| GCA_009619295 | WJOP00000000 | SAMN13181021 | 1244-C1-H3-B22 | Calditrichaeota | 2,223,554 | 459 | 2191 | 6,605 | 57 | 81 | 10 | 27 |
| GCA_009619035 | WJPB00000000 | SAMN13181033 | 1244-E5-H5-B19 | Spirochaeta | 1,522,441 | 390 | 1452 | 4,221 | 47 | 41 | 0 | 0 |
| GCA_009619125 | WJON00000000 | SAMN13181019 | 1244-C1-H3-B19 | Unclassified bacterium | 1,724,327 | 226 | 1686 | 9,503 | 53 | 61 | 2 | 0 |
| GCA_009619195 | WJOZ00000000 | SAMN13181031 | 1244-C3-H4-B23 | | 1,437,950 | 349 | 1342 | 4,977 | 41 | 59 | 1 | 0 |
| GCA_009618485 | WJOM00000000 | SAMN13181018 | 1244-C1-H3-B37 | | 1,464,732 | 418 | 1426 | 3,969 | 47 | 50 | 0 | 0 |
| GCA_009619155 | WJOW00000000 | SAMN13181028 | 1244-C3-H4-B1 | Hadesarchaea | 1,294,504 | 20 | 1460 | 148,197 | 50 | 88 | 4 | 17 |
| GCA_009618475 | WJOV00000000 | SAMN13181027 | 1244-F3-H4-B6 | Methanosarcinales (Euryarchaeota) | 1,093,430 | 165 | 1299 | 8,522 | 43 | 60 | 2 | 0 |
| GCA_009619135 | WJPD00000000 | SAMN13181035 | 1244-E5-H5-B17 | Unclassified archaeon | 1,307,191 | 249 | 1467 | 6,784 | 58 | 49 | 1 | 0 |
| GCA_009619335 | WJOS00000000 | SAMN13181024 | 1244-F2-H4-B13 | | 825,629 | 212 | 930 | 4,508 | 42 | 39 | 1 | 0 |

**Table S4. Matrix of average amino acid identities for the JS-1 Genus-1 MAGs.**

| | 35_14 | HGW-1 | E44_bin65 | MT.SAG.1 | UBA7920 | E10H5-B2 | 4572_76 |
|---|---|---|---|---|---|---|---|
| **35_14** | | | | | | | |
| **HGW-1** | 78 | | | | | | |
| **E44_bin65** | 78 | 74 | | | | | |
| **MT.SAG.1** | 76 | 74 | 77 | | | | |
| **UBA7920** | 81 | 79 | 79 | 78 | | | |
| **E10H5-B2** | 77 | 73 | 81 | 76 | 78 | | |
| **4572_76** | 78 | 76 | 72 | 73 | 80 | 76 | |
| **UBA9311** | 79 | 79 | 77 | 75 | 83 | 80 | 83 |

**Table S5. Hun-encoding genes in *Atribacteria* MAG B2 with percent identity to top hits in other MAGs.** Hun genes were also present in other MAGs from Site 1244 including *Firmicutes* E5B5-B3 (contigs 32 and 225).

| **Gene** | **PFAM** | **B2 contig E10H5_** | **B2 gene** | ***Atribacteria* bacterium isolate UBA7920** | ***Candidatus* Atribacteria bacterium MT.SAG.1** | ***Actinobacteria* RBG_13_35_12** | ***Atribacteria* JGI 0000014-F07** | ***Atribacteria* CG2_30_33_13** | ***Omnitrophica* WOR_2 SM23_29** | ***Omnitrophica* RBG_13_46_9** | ***Omnitrophica* RIFCSPHIGHO2 02_FULL_45_28** |
|---|---|---|---|---|---|---|---|---|---|---|---|
| ***hunR*** | PF02629 (CoA_binding), PF06971 (Put_DNA-bind_N) | C341 | RXG63122 | N/A | TFB08687 (88%) | OFW53501 (92%) | WP_09059864 (89%) | N/A | N/A | N/A | N/A |
| ***hunA*** | PF17179 (Fer4_22) | C341 | RXG63123 | HAJ31991 (87%) | TFB08688 (93%) | OFW53497 (91%) | WP_090598643 (89%) | OIP6690 (85%) | KPK38155 (68%) | OGW74721 (62%) | OGW81410 (63%) |
| ***hunB*** | PF00175 (NAD_binding_1), PF00970 (FAD_binding_6), PF10418 (DHODB_Fe-S_bind) | C341 | RXG63124 | HAJ31992 (94%) | TFB08716 (92%) | OFW53500 (91%) | WP_090598656 (93%) | OIP66908 (89%) | KPK38154 (77%) | OGW74722 (72%) | OGW81411 (69%) |
| ***hunC*** | PF13247 (Fer4_11) | C341 | RXG63125 | HAJ31993 (94%) | TFB08689 (93%) | OFW53499 (95%) | WP_090598644 (94%) | OIP66907 (91%) | KPK38158 (80%) | OGW74723 (78%) | OGW81412 (68%) |
| ***hunD*** | PF00146 (NADHdh) | C341 | RXG63126 | HAJ31994 (96%) | TFB08690 (95%) | OFW53496 (93%) | WP_090598646 (95%) | OIP66899 (92%) | KPK38153 (81%) | OGW74724 (79%) | OGW81413 (70%) |
| ***hunE*** | PF01058 (Oxidored_q6) | C341 | RXG63127 | HAJ31995 (98%) | TFB08691 (92%) | OFW53495 (96%) | WP_090598648 (95%) | OIP66906 (95%) | KPK39520 (90%) | OGW74725 (89%) | OGW81414 (85%) |
| ***hunF*** | PF00329 (Complex1_30kDa) | C341 | RXG63128 | HAJ31996 (87%) | TFB08692 (91%) | OFW53494 (85%) | WP_090598650 (83%) | OIP66898 (87%) | KPK39524 (73%) | OGW74727 (73%) | OGW81415 (68%) |
| ***hunG*** | PF00346 (Complex1_49kDa), PF00374 (NiFeSe_Hases) | C341 | RXG63129 | HAJ31997 (95%) | TFB08717 (96%) | OFW53493 (97%) | WP_090598653 (94%) | OIP66897 (94%) | KPK39519 (85%) | OGW74726 (87%) | OGW81416 (78%) |
| ***hunH*** | PF00361 (proton_antipo_M) | C341 | RXG63130 | HAJ31998 (94%) | TFB08693 (91%) | OFW53492 (88%) | WP_090599466 (88%) | OIP66896 (89%) | N/A | N/A | OGW81417 (77%) |
| ***hunI*** | PF13244 (DUF4040) | C1989 | RXG66691 | HAJ31999 (92%) | TFB08694 (95%) | OFW53491 (89%) | WP_090599464 (89%) | OIP66895 (80%) | KPK39517 (83%) | OGW74728 (81%) | OGW81418 (72%) |
| ***hunJ*** | PF04039 (MnhB) | C1989 | RXG66692 | HAJ32000 (97%) | TFB08695 (94%) | OFW53490 (95%) | WP_090599461 (85%) | OIP66894 (88%) | KPK40785 (85%) | OGW74729 (84%) | OGW81419 (80%) |
| ***hunK*** | PF00420 (Oxidored_q2) | C1989 | RXG66693 | HAJ32001 (96%) | TFB08696 (96%) | OFW53489 (91%) | WP_090599460 (90%) | OIP66893 (90%) | KPK40786 (80%) | N/A | OGW81420 (76%) |
| ***hunL*** | PF00361 (proton_antipo_M), PF00662 (proton_antipo_N), | C1989 | RXG66694 | HAJ32002 (96%) | TFB08697 (94%) | OFW53498 (87%) | WP_090599470 (82%) | OIP66892 (86%) | N/A | OGW74730 (76%) | OGW81421 (74%) |
| ***hunM*** | PF01899 (MNHE) | C1989 | RXG66695 | HAJ32003 (95%) | TFB08698 (98%) | OFW53488 (87%) | WP_090599458 (86%) | OIP66891 (86%) | KPK40787 (73%) | OGW74758 (73%) | OGW81422 (65%) |
| ***hunN*** | PF04066 (MrpF_PhaF) | C1989 | RXG66696 | HAJ32004 (96%) | TFB08699 (97%) | N/A | WP_090599455 (95%) | OIP66905 (91%) | KPK40788 (85%) | OGW74759 (85%) | OGW81430 (70%) |
| ***hunO*** | PF03334 (PhaG_MnhF_YufB) | C5669 | RXG63005 | HAJ32005 (91%) | TFB08700 (94%) | OFW53487 (91%) | WP_090599453 (88%) | OIP66890 (91%) | KPK40789 (77%) | OGW74731 (74%) | OGW81423 (70%) |
| ***hunP*** | PF12838 (Fer4_7) | C5669 | RXG63006 | HAJ32006 (89%) | TFB08701 (95%) | N/A | WP_090599451 (91%) | OIP66889 (88%) | KPK40790 (81%) | OGW74732 (76%) | OGW81424 (75%) |

**Table S6. Genes encoded on contigs containing the HTH-XRE transcriptional regulator/antitoxin AtiR in E10H5-B2 *Atribacteria* genomic bin, as shown in Figure 5, and percent identity to homologous genes in other (meta)genomes (excluding Hydrate Ridge hits).** Bolded genes were expressed in the metaproteome. *truncated protein; **contain conserved arginine residue for carboxylate transport.

| Annotation | Gene | Contig | Gene | Top hit (% identity) | Top *Atribacteria* hit in environmental genome or top hit in non-Hydrate Ridge genome |
|---|---|---|---|---|---|
| HTH-XRE regulator | *atiR* | E10H5_C107 | RXG62479 | HDK27529 (69%) | *Atribacteria* HyVt-22 |
| | | E10H5_C473 | RXG64192 | HDK27529 (66%) | *Atribacteria* HyVt-22 |
| | | E10H5_C194 | RXG65323 | HDK27529 (64%) | *Atribacteria* HyVt-22 |
| | | E10H5_C1742 | RXG64293 | HDK27529 (72%) | *Atribacteria* HyVt-22 |
| | | E10H5_C95 | RXG62928* | HDK27529 (65%) | *Atribacteria* HyVt-22 |
| | | E10H5_C238 | RXG62729 | HDK27529 (65%) | *Atribacteria* HyVt-22 |
| | | E10H5_C2 | RXG66887 | HDK27529 (63%) | *Atribacteria* HyVt-22 |
| | | E10H5_C2 | RXG66902 | HDK27529 (65%) | *Atribacteria* HyVt-22 |
| | | E10H5_C486 | RXG66788 | HDK27529 (59%) | *Atribacteria* HyVt-22 |
| | | E10H5_C687 | RXG66393 | HDK27529 (73%) | *Atribacteria* HyVt-22 |
| | | E10H5_C103 | RXG62795 | HDK27529 (66%) | *Atribacteria* HyVt-22 |
| | | E10H5_C85 | RXG66363 | HDK27529 (64%) | *Atribacteria* HyVt-22 |
| | | E10H5_C4712 | RXG63519 | HDK27529 (67%) | *Atribacteria* HyVt-22 |
| | | E10H5_C1316 | RXG65641 | HDK27529 (72%) | *Atribacteria* HyVt-22 |
| | | E10H5_C5280 | RXG63000 | HDK27529 (78%) | *Atribacteria* HyVt-22 |
| | | E10H5_C2922 | RXG65842 | HDK27529 (74%) | *Atribacteria* HyVt-22 |
| Tripartite tricarboxylate transporter | *tctC* | E10H5_C95 | RXG62929 | WP_066240574 (53%) | *Anaerosporomusa subterranea* |
| | *tctB* | | RXG62930 | WP_118245178 (35%) | *Clostridium* AM58-1XD |
| | *tctA* | | RXG62931 | WP_093692001 (54%) | *Sporolituus thermophilus* |
| | *tctC* | E10H5_C107 | RXG62483 | MBN2395378 (71%) | *Atribacteria* Zod_Metabat.864 |
| | *tctB* | | RXG62484 | MBN2395379 (53%) | *Atribacteria* Zod_Metabat.864 |
| | *tctA* | | RXG62485 | MBN2395380 (68%) | *Atribacteria* Zod_Metabat.864 |
| | *sat* | | RXG62486 | MBN2395381 (68%) | *Atribacteria* Zod_Metabat.864 |
| | *hyp* | | RXG62487 | MBN2395382 (61%) | *Atribacteria* Zod_Metabat.864 |
| C4-dicarboxylate transporter | *dctP* | E10H5_C95 | RXG62935** | PKL21426 (55%) | *Spirochaetae* HGW-1 |
| | ***dctQ*** | | **RXG62936** | **AEG13811 (34%)** | ***Desulfofundulus kuznetsovii* DSM 6115** |
| | *dctM* | | RXG62937 | PKL21182 (57%) | *Spirochaetae* HGW-4 |
| | *dctP* | E10H5_C238 | RXG62728 | HBY57297 (84%) | *Atribacteria* UBA9904 |
| | *dctQ* | | RXG62727 | HBY57298 (74%) | *Atribacteria* UBA9904 |
| | *dctM* | | RXG62726 | HBY57299 (80%) | *Atribacteria* UBA9904 |
| | *dctP* | E10H5_C486 | RXG66786** | HDK26676 (86%) | *Atribacteria* HyVt-22 |
| | *dctQ* | | RXG66785 | HDK26675 (69%) | *Atribacteria* HyVt-22 |
| | *dctM* | | RXG66784 | HDK26674 (82%) | *Atribacteria* HyVt-22 |
| | *dctP* | E10H5_C664 | RXG63168 | HAJ33545 (72%) | *Atribacteria* UBA7920 |
| | *dctQ* | | RXG63170 | HAJ33544 (73%) | *Atribacteria* UBA7920 |
| | *dctM* | | RXG63171 | HAJ33543 (84%) | *Atribacteria* UBA7920 |
| Glycerol-3-phosphate transporter | *ugpB* | E10H5_C85 | RXG66368 | KPJ85459 (68%) | *Spirochaetes* DG_61 |
| | *ugpA* | | RXG66369 | KPJ85458 (67%) | *Spirochaetes* DG_61 |
| | *ugpE* | | RXG66370 | KPJ85457 (71%) | *Spirochaetes* DG_61 |
| | *ugpB* | E10H5_C103 | RXG62790 | WP_068137292 (47%) | *Limnochorda pilosa* |

| | | | | | |
|---|---|---|---|---|---|
| | *ugpA* | | RXG62789 | WP_082726097 (46%) | *Limnochorda pilosa* |
| | *ugpE* | | RXG62788 | WP_068137283 (46%) | *Limnochorda pilosa* |
| Branched chain | *livH* | E10H5_C2 | RXG66901 | PKP61938 (90%) | *Atribacteria* HGW-1 |
| amino acid | *livM* | | RXG66900 | PKP61939 (85%) | *Atribacteria* HGW-1 |
| transporters | *livG* | | RXG66899 | PKP61948 (85%) | *Atribacteria* HGW-1 |
| | *livF* | | RXG66898 | PKP61940 (80%) | *Atribacteria* HGW-1 |
| | *livK* | | RXG66897 | PKP61941 (89%) | *Atribacteria* HGW-1 |
| | *livH* | E10H5_C238 | RXG62732 | RLE68614 (60%) | *Thermoprotei* archaeon |
| | *livM* | | RXG62736 | RLE68615 (55%) | *Thermoprotei* archaeon |
| | *livG* | | RXG62731 | RLE68616 (57%) | *Thermoprotei* archaeon |
| | *livF* | | RXG62730 | RLE68617 (56%) | *Thermoprotei* archaeon |
| | *livK* | E10H5_C687 | RXG66394 | OQY40502 (43%) | *Atribacteria* 4572_76 |
| | *livH* | | RXG66395 | OQY40503 (41%) | *Atribacteria* 4572_76 |
| | *livM* | | RXG66396 | OQY40504 (42%) | *Atribacteria* 4572_76 |
| | *livG* | | RXG66397 | OGP70799 (50%) | *Deltaproteobacteria* RBG_16_50_11 |
| | *livF* | | RXG66398 | OQY40505 (55%) | *Atribacteria* 4572_76 |
| | ***livH*** | **E10H5_C8009** | **30420** | **OQY40503 (95%)** | ***Atribacteria* 4572_76** |
| | *livM* | | 30421 | OQY40504 (93%) | *Atribacteria* 4572_76 |
| Purine transporter | ***bmpA*** | **E10H5_C473** | **RXG64193** | **PKP58720 (94%)** | ***Atribacteria* HGW-1** |
| | | | RXG64197 | PKP60518 (88%) | *Atribacteria* HGW-1 |
| | | | RXG64194 | PKP60517 (94%) | *Atribacteria* HGW-1 |
| | | | RXG64195 | PKP60516 (96%) | *Atribacteria* HGW-1 |
| M20 | | **E10H5_C194** | RXG65322 | WP_034420537 (72%) | *Clostridiales* DRI-13 |
| (zinc peptidase, | | | RXG65321 | WP_034420536 (73%) | *Clostridiales* DRI-13 |
| amidohydrolase) | | | RXG65329 | WP_034420535 (75%) | *Clostridiales* DRI-13 |
| Sulfur-related | *sat* | E10H5_C107 | RXG62486 | HDK26586 (85%) | *Atribacteria* HyVt-22 |
| (and neighboring) | | | RXG62487 | HDK26585 (67%) | *Atribacteria* HyVt-22 |
| Other genes | *gabT* | E10H5_C4712 | RXG63518 | PKP55816 (87%) | *Atribacteria* HGW-1 |
| | C69 | E10H5_C2795 | RXG64595 | WP_093794159 (43%) | *Sporomusa acidovorans* |
| | RTX | E10H5_C1316 | RXG65639 | OGD35967 (42%) | *Atribacteria* RBG_16_35_8 |
| | RTX/Ig | E10H5_C2922 | RXG65844 | OGD35967 (43%) | *Atribacteria* RBG_16_35_8 |

36

**Table S7. Putative toxin-antitoxin systems in *Atribacteria* E10H5-B2, and percent identity to homologous genes in other genomes.**

| Annotation | Gene | Contig | Gene | Top hit (% identity) | Top hit |
|---|---|---|---|---|---|
| Antitoxin | MazE-like | E10H5_ C26 | RXG62841 | PIU28774 (82%) | *Actinobacteria* CG08_land_8_20_14_0_20_35_9 |
| Toxin | MazF-like | | RXG62840 | PIU28775 (88%) | *Actinobacteria* CG08_land_8_20_14_0_20_35_9 |
| Antitoxin | NikR-like | E10H5_C33 | RXG63986 | PIU25819 (95%) | *Atribacteria* CG08_land_8_20_14_0_20_33_29 |
| Toxin | MazF-like | | RXG63985 | PIU25820 (83%) | *Atribacteria* CG08_land_8_20_14_0_20_33_29 |
| Antitoxin | MazE-like | E10H5_C68 | RXG64905 | PKP54844 (89%) | *Atribacteria* HGW-1 |
| Toxin | MazF-like | | RXG64906 | PKP54843 (84%) | *Atribacteria* HGW-1 |
| Antitoxin | VapC-like | E10H5_C81 | RXG64104 | PKP54729 (90%) | *Atribacteria* HGW-1 |
| Toxin | MazE-like | | RXG64105 | PKP54728 (81%) | *Atribacteria* HGW-1 |
| Antitoxin | MazE-like | E10H5_C81 | RXG64330 | PIY33044 (92%) | *Atribacteria* CG_4_10_14_3_um_filter_34_13 |
| Toxin | MazF-like | | RXG64331 | PIY33045 (92%) | *Atribacteria* CG_4_10_14_3_um_filter_34_13 |
| Antitoxin | MazE-like | E10H5_C184 | RXG64099 | OQY39409 (97%) | *Atribacteria* 4572_76 |
| Toxin | MazF-like | | RXG64098 | OQY39408 (98%) | *Atribacteria* 4572_76 |
| Antitoxin | MazE-like | E10H5_C1146 | RXG64860 | WP_090599534 (91%) | *Atribacteria* JGI 0000014-F07 |
| Toxin | MazF-like | | RXG64859 | WP_090599536 (89%) | *Atribacteria* JGI 0000014-F07 |
| Antitoxin | MazE-like | E10H5_C1351 | RXG65145 | PKP54770 (94%) | *Atribacteria* HGW-1 |
| Toxin | MazF-like | | RXG65146 | PKP54769 (89%) | *Atribacteria* HGW-1 |
| Antitoxin | MazE-like | E10H5_ C5010 | RXG67035 | WP_078128337 (59%) | *Leptospira alexanderi* |
| Toxin | MazF-like | | RXG67036 | WP_010577027 (74%) | *Leptospira alexanderi* |

**Table S8. Glycosyltransferase contigs in E10-H5-B2 *Atribacteria* genomic bin, and percent identity to homologous genes in other genomes.**

| Annotation | Contig | Gene | Top hit (% identity) | Top hit |
|---|---|---|---|---|
| Glycosyltransferase 2<br>Glycosyltransferase 1<br>Glycosyltransferase 2<br>Glycosyltransferase 2<br>Glycosyltransferase 2<br>NAD-dependent epimerase/dehydratase<br>NDP-sugar synthase<br>GDP-mannose 4,6-dehydratase | E10H5_C38 | N/A<br>RXG64781<br>RXG64782<br>RXG64783<br>RXG64791<br>RXG64792<br>RXG64793<br>RXG64794 | OFW63513 (72%)<br>OFW63514 (72%)<br>OGD31966 (66%)<br>WP_104083790 (34%)<br>GBC98638 (43%)<br>OFW55242 (82%)<br>OFW55241 (82%)<br>OFW55240 (88%) | *Actinobacteria* RBG_19FT_COMBO_36_27<br>*Actinobacteria* RBG_19FT_COMBO_36_27<br>*Atribacteria* RBG_16_35_8<br>*Cryobacterium* Y11<br>Bacterium HR17<br>*Actinobacteria* RBG_13_35_12<br>*Actinobacteria* RBG_13_35_12<br>*Actinobacteria* RBG_13_35_12 |
| Glycosyltransferase 1<br>Glycosyltransferase 1<br>Glycosyltransferase 1<br>Glycosyltransferase 1<br>O-antigen ligase<br>Glycosyltransferase 1 | E10H5_C81 | RXG64335<br>RXG64336<br>RXG64337<br>RXG64338<br>RXG64339<br>RXG64340 | WP_093394743 (45%)<br>WP_093394743 (53%)<br>OGI06781 (40%)<br>SPE32180 (46%)<br>OGD19921 (30%)<br>OGD19922 (55%) | *Thermodesulforhabdus norvegica*<br>*Thermodesulforhabdus norvegica*<br>*Melainabacteria* RIFCSPLOWO2_12_FULL_35_11<br>*Solibacteres* SbA2<br>*Aminicenantes* RBG_13_64_14<br>*Aminicenantes* RBG_13_64_14 |
| Polysaccharide (de)acetylase<br>Exopolysaccharide polyprenyl<br>glycosylphosphotransferase<br>Sugar epimerase<br>Glycosyltransferase 1 (Cap1E-like)<br>UDP-N-acetylglucosamine 4-epimerase<br>Vi polysaccharide biosynthesis protein<br>Phospholipid carrier-dependent<br>glycosyltransferase<br>Glycosyltransferase<br>Phospholipid carrier-dependent<br>glycosyltransferase | E10H5_C230 | RXG63410<br>RXG63411<br>RXG63412<br>RXG63413<br>RXG63414<br>RXG63415<br>RXG63416<br>RXG63417<br>RXG63418 | WP_019599125 (28%)<br>PKP61696 (88%)<br>PKP61697 (77%)<br>WP_071120025 (57%)<br>WP_036938033 (71%)<br>WP_071120026 (68%)<br>WP_036935155 (48%)<br>PJE73714 (51%)<br>WP_036935155 (43%) | *Rhodonellum* spp.<br>*Atribacteria* HGW-1<br>*Atribacteria* HGW-1<br>*Romboutsia timonensis*<br>*Pseudobacteroides cellulosolvens*<br>*Romboutsia timonensis*<br>*Pseudobacteroides cellulosolvens*<br>Terrybacteria CG10_big_fil_rev_8_21_14_0_10_41_10<br>*Pseudobacteroides cellulosolvens* |
| Glycosyltransferase 1<br>GDP-fucose synthetase<br>GDP-mannose 4,6-dehydratase | E10H5_C3323 | RXG63206<br>RXG63207<br>(2,613-2,990) | PKP61698 (81%)<br>PKP61699 (85%)<br>PKP61700 (82%) | *Atribacteria* HGW-1<br>*Atribacteria* HGW-1<br>*Atribacteria* HGW-1 |
| UDP-glucose 6-dehydrogenase (Ugd)<br>NAD-dependent epimerase/dehydratase<br>Aminotransferase<br>Formyltransferase (non-ribosomal<br>peptide synthetase-like)<br>Acetyltransferase<br>Deacetylase | E10H5_C266 | RXG63573<br>RXG63566<br>RXG63567<br>RXG63568<br>RXG63569<br>RXG63570 | PKP61722 (77%)<br>PKP59007 (93%)<br>PKP59008 (93%)<br>PKP59009 (90%)<br>PKP59010 (87%)<br>RME50047 (61%) | *Atribacteria* HGW-1<br>*Atribacteria* HGW-1<br>*Atribacteria* HGW-1<br>*Atribacteria* HGW-1<br>*Atribacteria* HGW-1<br>*Deltaproteobacteria* bacterium |
| 7-keto-8-aminopelargonate synthetase<br>UDP-glucose 4-epimerase | E10H5_C306 | RXG65589<br>RXG65590 | OGD15025 (91%)<br>OGD15024 (84%) | *Atribacteria* RBG_19FT_COMBO_35_14<br>*Atribacteria* RBG_19FT_COMBO_35_14 |
| UDP-glucose 4-epimerase | E10H5_C441 | RXG65590 | OGD15024 (87%) | *Atribacteria* RBG_19FT_COMBO_35_14 |
| Radical SAM P-methyltransferase | E10H5_C558 | RXG63508 | OGD16377 (81%) | *Atribacteria* RBG_19FT_COMBO_35_14 |
| Glycosyltransferase 1 | E10H5_C1163 | RXG62499 | WP_090599978 (92%) | *Atribacteria* JGI 0000014-F07 |
| Glycosyltransferase 1 (Cap1E-like)<br>Exopolysaccharide polyprenyl<br>glycosylphosphotransferase<br>Acetyltransferase<br>Polysaccharide biosynthesis protein | E10H5_C3994 | RXG63236<br>RXG63237<br>RXG63239<br>RXG63238 | WP_071605078 (53%)<br>PKP62045 (76%)<br>PKP62044 (84%)<br>PKP62043 (87%) | *Anaeroarcus burkinensis*<br>*Atribacteria* HGW-1<br>*Atribacteria* HGW-1<br>*Atribacteria* HGW-1 |

| | | | | |
|---|---|---|---|---|
| Glycosyltransferase 4<br>Cyclic nucleotide-binding domain-containing protein | E10H5_C4124 | RXG64087<br>RXG64088 | PKP55791 (95%)<br>PKP55792 (93%) | *Atribacteria* HGW-1<br>*Atribacteria* HGW-1 |
| NAD-dependent epimerase/dehydratase<br>Glycosyltransferase 4 | E10H5_C4631 | RXG66927<br>RXG66928 | PKP58948 (97%)<br>PKP58071 (83%) | *Atribacteria* HGW-1<br>*Atribacteria* HGW-1 |
| Bacillithiol biosynthesis deacetylase<br>Glycosyltransferase 2 | E10H5_C4676 | RXG65226<br>RXG65225 | OGD13744 (93%)<br>OGW00676 (55%) | *Atribacteria* RBG_19FT_COMBO_35_14<br>*Nitrospinae* RIFCSPLOWO2_01_FULL_39_10 |

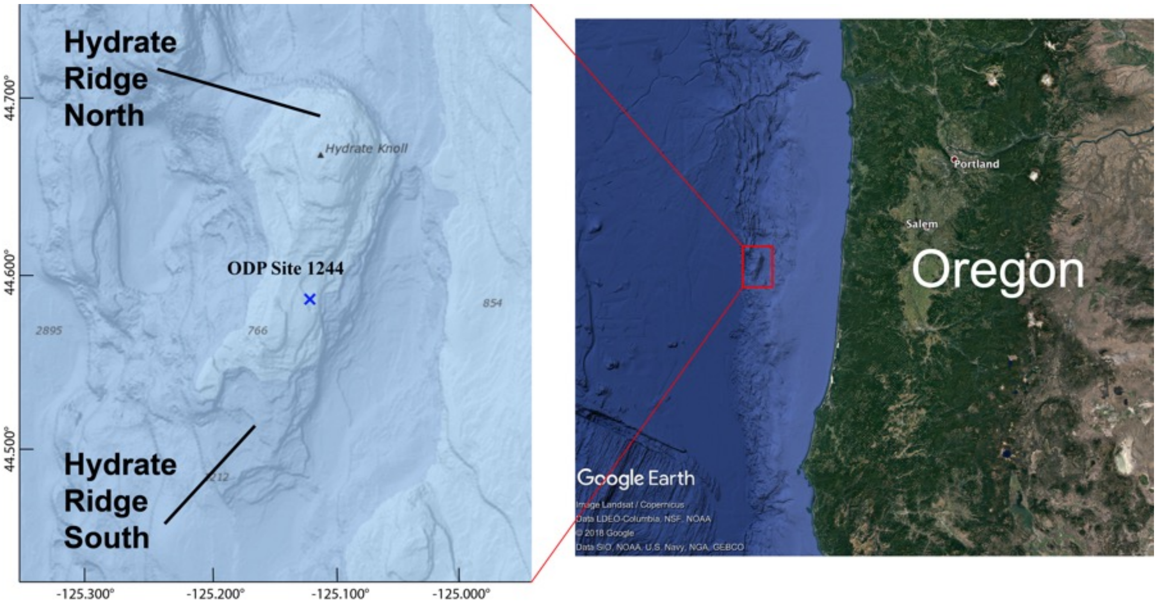


**Figure S1.** Map of Site ODP 1244 (44°35.1784´N; 125°7.1902´W) on Hydrate Ridge, drilled on IODP Leg 204. The site is located 80 km west of Oregon, in the accretionary complex of the Cascadia subduction zone, on the eastern flank of Hydrate Ridge, ~3 km northeast of the southern summit.

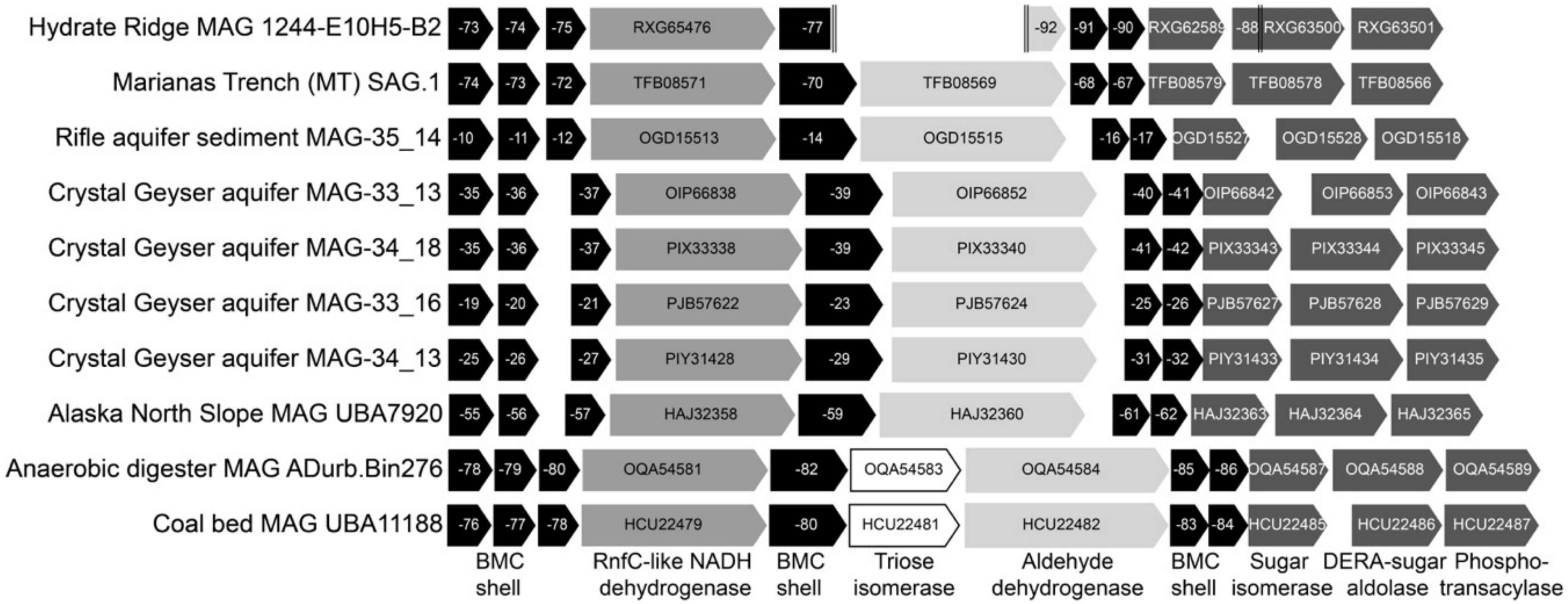


**Figure S2.** Bacterial microcompartment (BMC) gene loci from *Atribacteria* (JS-1) MAGs and SAGs in UniProt. Each gene is labeled with its NCBI accession. Short genes are labeled with the last two digits of the NCBI accession (see neighboring genes for remainder of accession). The BMC loci from Hydrate Ridge MAG 1244-E10H5-B2 is distributed over three short contigs with some genes truncated (indicated by double vertical lines).
